# Using artificial intelligence to document the hidden RNA virosphere

**DOI:** 10.1101/2023.04.18.537342

**Authors:** Xin Hou, Yong He, Pan Fang, Shi-Qiang Mei, Zan Xu, Wei-Chen Wu, Jun-Hua Tian, Shun Zhang, Zhen-Yu Zeng, Qin-Yu Gou, Gen-Yang Xin, Shi-Jia Le, Yin-Yue Xia, Yu-Lan Zhou, Feng-Ming Hui, Yuan-Fei Pan, John-Sebastian Eden, Zhao-Hui Yang, Chong Han, Yue-Long Shu, Deyin Guo, Jun Li, Edward C. Holmes, Zhao-Rong Li, Mang Shi

## Abstract

Current metagenomic tools can fail to identify highly divergent RNA viruses. We developed a deep learning algorithm, termed LucaProt, to discover highly divergent RNA-dependent RNA polymerase (RdRP) sequences in 10,487 metatranscriptomes generated from diverse global ecosystems. LucaProt integrates both sequence and predicted structural information, enabling the accurate detection of RdRP sequences. Using this approach we identified 161,979 potential RNA virus species and 180 RNA virus supergroups, including many previously poorly studied groups, as well as RNA virus genomes of exceptional length (up to 47,250 nucleotides) and genomic complexity. A subset of these novel RNA viruses were confirmed by RT-PCR and RNA/DNA sequencing. Newly discovered RNA viruses were present in diverse environments, including air, hot springs and hydrothermal vents, and both virus diversity and abundance varied substantially among ecosystems. This study advances virus discovery, highlights the scale of the virosphere, and provides computational tools to better document the global RNA virome.

**In brief:** A deep learning algorithm (LucaProt) that integrates both sequence and predicted structural information was employed to identify highly divergent RNA viral “dark matter” in 10,487 metatranscriptomes from diverse global ecosystems. A total of 161,979 potential RNA virus species and 180 RNA virus supergroups was unveiled using this AI approach, including many understudied groups.

## Introduction

RNA viruses infect a diverse array of host species. Despite their omnipresence, the pivotal role of RNA viruses as major constituents of global ecosystems has only recently garnered recognition due to large-scale virus discovery initiatives in animals,^1,2^ plants,^3^ fungi,^4^ aquatic environments,^5^ marine,^6^ soil environments,^7^ and planetary metatranscriptomes.^8^ A common characteristic of these studies is their reliance on the analysis of RNA-dependent RNA polymerase (RdRP) sequences, a canonical component of RNA virus genomes. Collectively, these studies have led to the identification of tens of thousands of novel virus species, resulting in at least a 10-fold expansion of the virosphere and the proposal of new phylum-level virus groups such as the “*Taraviricota*” (i.e., “quenyaviruses”).^6,9^ Similarly, the data mining of metatranscriptomes from diverse ecosystems has revealed several divergent clades of RNA bacteriophage,^10,11^ while recent metatranscriptomic studies have led to a remarkable 5-fold expansion in the diversity of viroid-like circular RNAs.^12–14^ Despite such progress in uncovering RNA virus diversity through ecological sampling and sequencing, it is probable that more divergent groups of RNA viruses remain to be discovered.^9,15^ This is in part because the current tools for metagenomic identification of RNA viruses can miss some highly divergent RdRPs.^16^ It is therefore imperative to develop innovative strategies for the efficient identification of the full spectrum of RNA virus diversity.

Over the past decade, artificial intelligence (AI) related approaches, especially deep learning algorithms, have had a major impact on various research fields in the life sciences, such as molecular docking, compound screening and interaction, protein structure prediction and functional annotation, and infectious disease modelling.^17–22^ This progress can be attributed to the advantages of deep learning algorithms over classic bioinformatic approaches, including enhanced accuracy, superior performance, reduced reliance on feature engineering, flexible model architectures, and self-learning capabilities.^23,24^ Recently, deep learning approaches, such as CHEER, VirHunter, Virtifier and RNN-VirSeeker have been developed and applied to the identification of viruses from genomic and metagenomic data.^25–28^

These tools employ convolutional neural networks (CNNs) and recurrent neural networks (RNNs). CNNs are specifically designed for processing spatial data such as images and leverage convolutions to exploit local correlations,^29^ whereas RNNs are adept at handling sequential data by capturing temporal dependencies and serial order memory.^30^ Despite their versatility, both face limitations in processing biological sequences: CNNs may encounter challenges with inputs of varying lengths and capturing global correlations, while RNNs struggle with longer sequences due to vanishing or exploding gradients and difficulties in capturing long-term dependencies. It is imperative to consider these shortcomings when assessing their appropriateness for specific tasks. In addition, many of these methodologies exclusively focus on nucleotide sequences, disregarding protein sequences or structural information, thereby constraining their capacity to identify highly divergent RNA viruses. Recently, the transformer architecture has emerged as a powerful alternative for protein function predictions based on sequence data, effectively accommodating sequences of varying lengths and efficiently capturing both local and long-range relationships across sequence positions, surpassing the capabilities of CNNs and RNNs.^31–34^ Consequently, the transformer architecture can be leveraged to design better tools for identifying highly divergent RNA viruses.

Herein, we present a transformer-based tool for RNA virus discovery that utilizes protein sequences and the structural characteristics of viral RdRP sequences. This tool was applied to a data set comprising 10,487 metatranscriptomes from diverse ecological systems. To validate and perform comparative analysis, the same data set was processed using other available bioinformatics tools, and 50 samples were analyzed using both DNA and RNA sequencing. By employing this tool in conjunction with extensive sequence data, we demonstrate how artificial intelligence can accurately and efficiently detect RNA viruses exhibiting genetic divergence beyond the capabilities of traditional similarity-based methods, revealing previously unrecognized viral diversity.

## Results

### Deep learning reveals the dark matter of the RNA virosphere

We performed systematic searches to expand the diversity of RNA viruses in a variety of ecological systems sampled on a global scale (Figures 1 and 2; Tables S1 and S2). Accordingly, a total of 10,487 metatranscriptomes (51 Tb of total sequencing data) were assembled, which resulted in more than 1,368 million contigs and 872 million predicted proteins. Based on this data set, potential viral RdRPs were revealed and cross-validated using two different strategies (Figures 1A, S1-2). The major AI algorithm used here, denoted “LucaProt”, is a deep learning, transformer-based model based on sequence and structural features of 5,979 well-characterized viral RdRPs (positive samples) and 229,434 protein sequences that were not viral RdRPs (negative samples), including non-RdRP viral proteins, reverse transcriptases, and cellular proteins. LucaProt demonstrated exceptional accuracy (0.014% false positives) and specificity (1.72% false negatives) when evaluated on the test data set (Figure 1B). Furthermore, the generalizability and robustness of LucaProt were further confirmed through a 10-fold cross-validation analysis (Figure S2). Independently to LucaProt, we applied a more conventional approach (i.e., “ClstrSearch”) that clustered all proteins based on their sequence similarity and then used BLAST or HMM models to identify resemblance to viral RdRPs or non-viral RdRP proteins.

**Figure 1.**
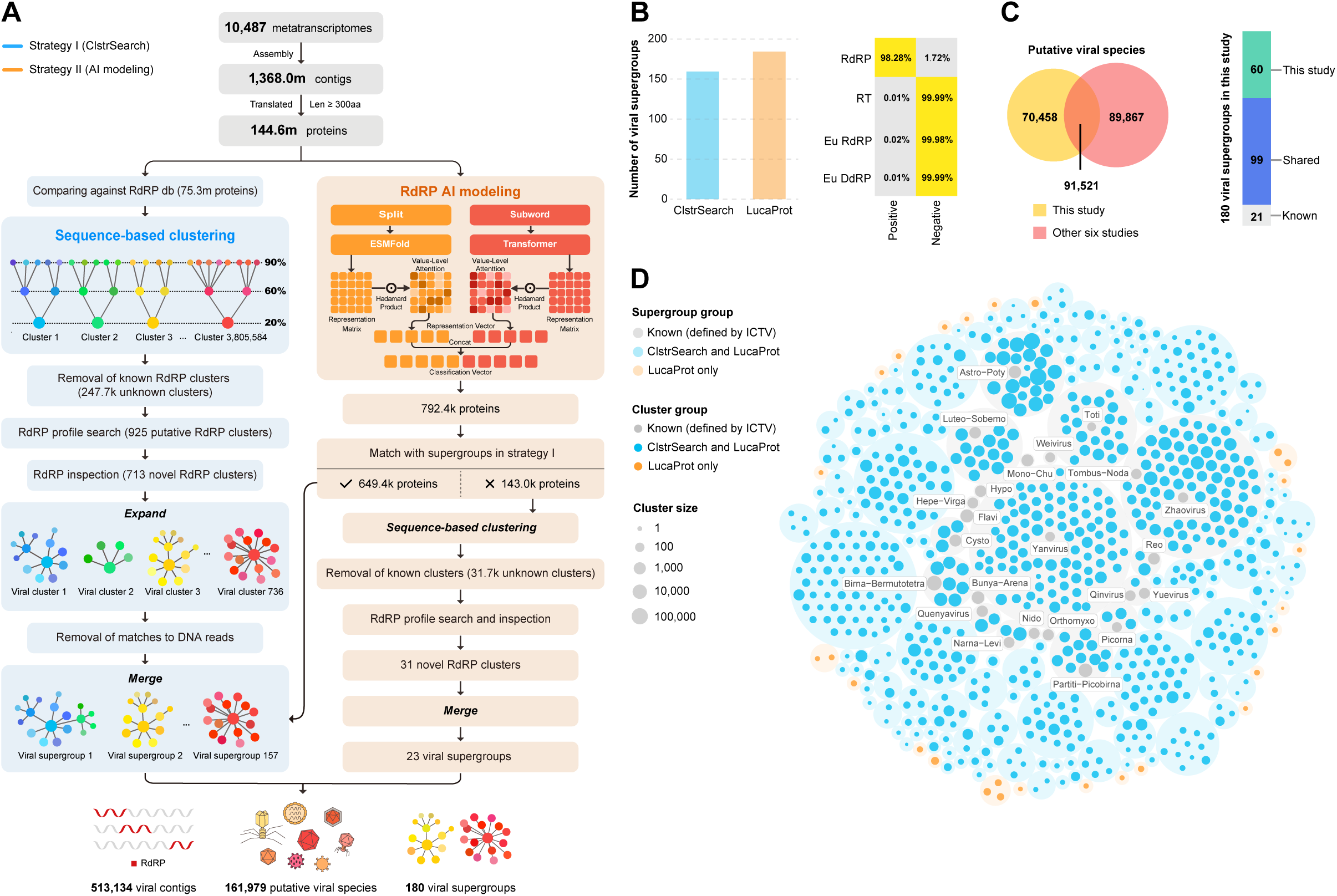
The global RNA virosphere. (A) RNA virus discovery pipeline. The pathway for sequence homolog-based virus discovery is highlighted in blue on the left, including the clustering, expand and merge steps. The RdRP AI modeling pathway is highlighted in orange on the right, including the modeling, clustering and merge steps. (B) Number of viral supergroups discovered using two methods (left), and the detection accuracy of LucaProt (right). (C) Venn diagram shows the shared putative virus species between available data from Wolf *et al*., Edgar *et al*., Zayed *et al*., Neri *et al*., Chen *et al*., Olendraite *et al*., and this study. The bar graph shows the shared viral supergroups between the seven studies and the unique viral supergroups identified in this study. (D) Diverse RNA virus clusters (dark colored small circle) and RNA virus supergroups (light colored large circle). The known viral clusters and supergroups defined by ICTV are shown in dark grey and light grey, respectively. The viral clusters and supergroups discovered by both ClstrSearch and LucaProt are shown in dark blue and light blue, respectively. The viral clusters and supergroups additionally discovered by LucaProt only are in shown dark orange and light orange, respectively. See also Figures S1-S3 and Tables S1-S3.

**Figure 2.**
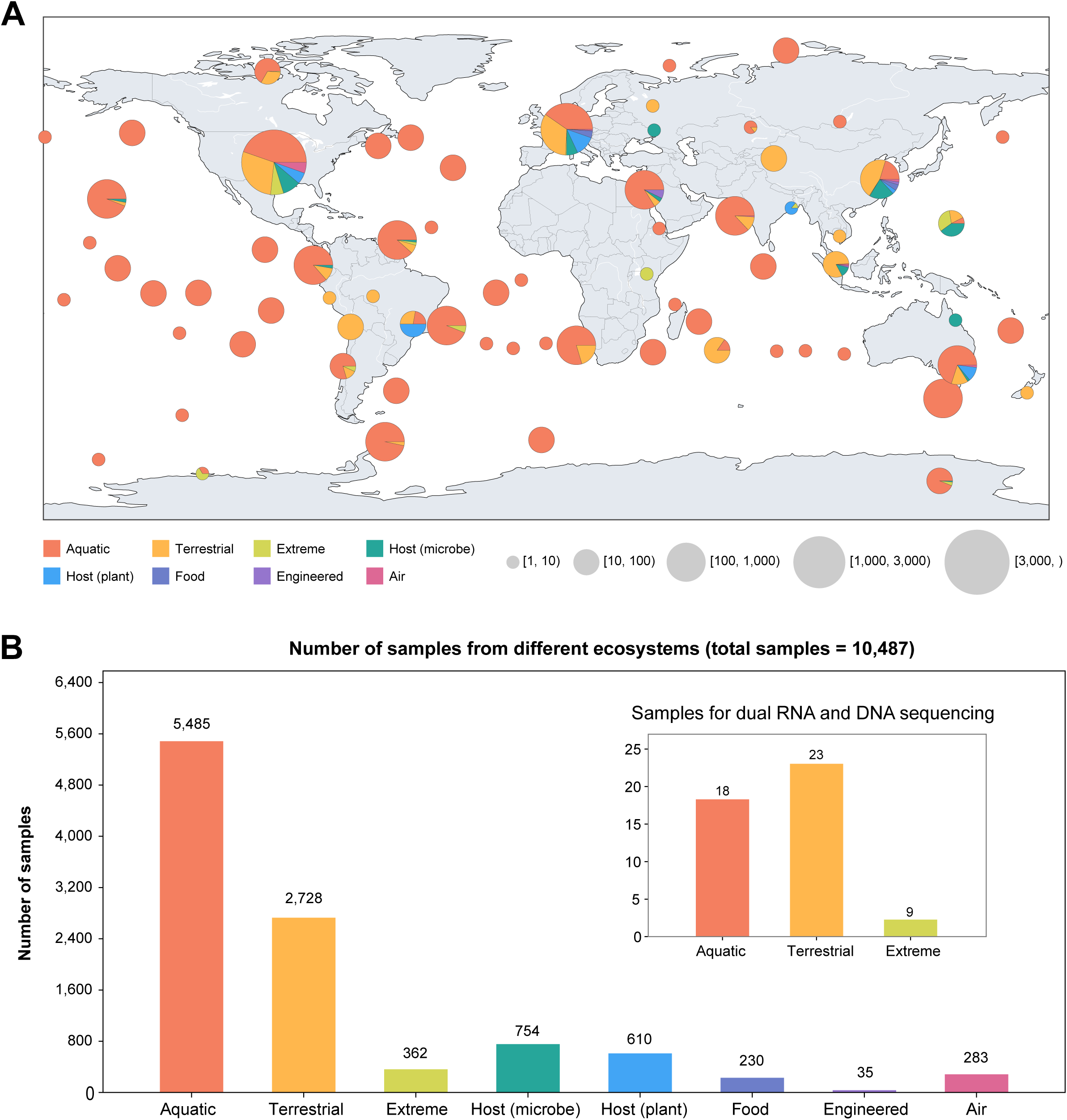
Geographic coverage of the metatranscriptomic data analyzed in this study. (A) Geographical distribution of the samples analyzed at an ecosystem level. Pie size is positively correlated to the number of samples (log10). The DBSCAN clustering algorithm was applied to group 1,837 latitude and longitude points from all metatranscriptomes into 70 clustered points. (B) Total number of samples at different ecosystems. The embedded bar chart represents the samples used for dual RNA and DNA sequencing in this study.

By merging the results of the two search strategies we discovered 513,134 RNA viral contigs, representing 161,979 putative viral species (i.e., >90% RdRP identity), and 180 RNA viral supergroups that are comparable to existing viral classes and phyla as defined by the International Committee on Taxonomy of Viruses (ICTV) (Figure 1; Table S3; see Methods). Subsequently, we performed an automatic comparison of RdRP protein sequences with a uniform definition (identity≥0.9, aligned fraction≥0.3) among this and other studies to reveal a total of 70,458 putative unique viral species newly identified by LucaProt (Figures 1C and S3). Notably, we unveiled 60 previously unidentified and underexplored groups that have received only limited attention to date^5–8,10,35^ (Figures 1C and S3). Among these, 512,690 viral contigs (99.9% of total contigs) and 157 supergroups (87.2%) were identified by both LucaProt and ClstrSearch, while an additional 444 contigs and 23 supergroups were only identified by LucaProt (Figure 1D).

### Benchmarking of LucaProt with other tools for virus discovery

To assess the sensitivity and specificity of LucaProt, we benchmarked it against four other virus discovery tools, utilizing the same data set and RdRP database. Notably, LucaProt exhibited the highest recall rate (i.e., proportion of correctly predicted true positives), while maintaining a relatively low false positive rate (i.e., proportion of incorrectly predicted true negatives as positives; Figure 3A), along with reasonable computational efficiency (Figure 3E). Specifically, in comparisons, HMMscan and PalmScan showed the highest precision rates (99.80%, 99.46%) but the lowest recall rates (65.00%, 65.03%), while Diamond blast and HH-suite both exhibited the highest false positive rate (0.195%, 0.573%; Figures 3A). When considering all RdRPs identified in this study, LucaProt exhibited the most comprehensive virus discovery (98.22%), whereas the other four virus discovery tools could only identify a portion of the entire collection (76.82%-87.81%, Figure 3B). Importantly, our benchmarking results demonstrated that the other virus discovery tools only identified a minority (<42%) of the new viruses identified by LucaProt only (i.e., but not by ClstrSearch) (Figure 3C).

**Figure 3.**
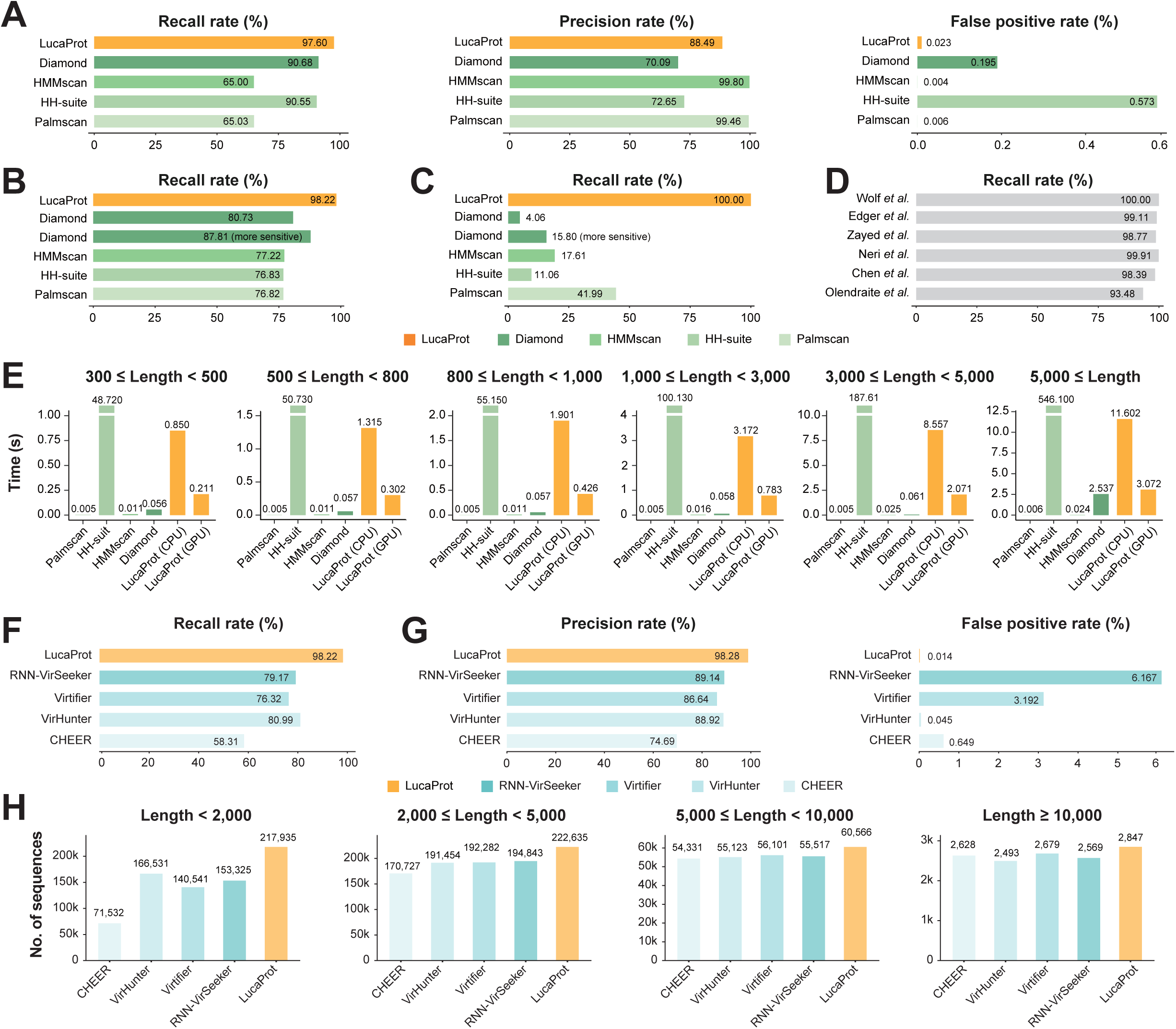
Benchmarking of LucaProt with other virus discovery tools. (A) The recall, precision and false positive rate of LucaProt, Diamond, HMMscan, HH-suite, and PalmScan was compared using the 50 metatranscriptomes sequenced in this study. (B) The recall of LucaProt, Diamond, HMMscan, HH-suite, and PalmScan in identifying all the RdRPs identified in this study. (C) The recall of LucaProt, Diamond, HMMscan, HH-suite, and PalmScan compared for the RdRPs additionally identified by LucaProt only. (D) Recall of LucaProt in identifying RdRPs (length≥300aa) from six previous studies. (E) The average time used by each bioinformatic tool was calculated based on six data sets of varying lengths, each comprising 50 positive sequences and 50 negative sequences. (F) Recall rate of prediction results for CHEER, VirHunter, Virtifier, RNN-VirSeeker and LucaProt based on all the RdRPs identified this study. (G) Precision rate and false positive rate of prediction results for CHEER, VirHunter, Virtifier, RNN-VirSeeker and LucaProt based on the test data set. (H) Number of viral sequences of different groups by contig length identified by CHEER, VirHunter, Virtifier, RNN-VirSeeker and LucaProt. The training machines, training data sets, training strategies, and final model selection of all methods compared are consistent with LucaProt. All comparisons utilized multiple sets of hyperparameters with the best results selected in each case.

Of note, LucaProt successfully recalled over 98% of the virus RdRPs identified by six other published studies,^5–8,10,35^ even though none were used in either training or testing of the models (Figure 3D). The exception was the study of Olendraite *et al.,*^35^ for which LucaProt had a slightly lower recall rate (Figure 3D). Manual inspection and sequence alignment revealed that the sequences from Olendraite *et al.,*^35^ not detected by LucaProt lacked the core RdRP domain region. LucaProt also outperformed CHEER, VirHunter, Virtifier and RNN-VirSeeker RNA virus discovery tools^25–28^ in terms of recall, precision and long sequence processing (Figures 3F-3H). The advanced transformer architecture incorporated into LucaProt allowed the parallel processing of longer amino acid sequences,^31,36^ which can better capture the relationships between residues from distant parts of sequence space than the CNN and/or RNN encoders implemented in the other bioinformatic tools compared.

### Verification and confirmation of newly identified viral supergroups

That the 180 viral supergroups identified here represented RNA rather than DNA sequences was substantiated through multiple lines of evidence. At the sequence level, two criteria were employed to establish a viral supergroup: the absence of similarity to cellular proteins and the presence of key RdRP motifs (Figures 4A and S4). Moreover, the majority (60/180) of the newly identified supergroups, including those from LucaProt, shared varying degrees of sequence similarity with existing reference RdRPs (i.e., BLAST *e*-value≤1*E*-3 and/or had HMM model score≥10) (Figures 4A and S4; Table S4). To validate computational predictions, simultaneous DNA and RNA extraction and sequencing were performed on the 50 environmental samples collected in this study to examine the presence of the 115 viral supergroups identified in these samples (Table S5). This revealed that only RNA sequencing reads mapped to contigs associated with viral RdRPs, whereas both RNA and DNA sequencing reads mapped to contigs linked with DNA viruses, reverse-transcriptase (RT), and cellular organisms (Figures 4B and S5). Seventeen of 115 viral supergroups were further confirmed by a more sensitive RT-PCR approach: this revealed an absence of sequences encoding viral RdRP in the DNA extractions, suggesting that these viral supergroups are *bona fide* RNA organisms (Figures 4C and S5B). Finally, a 3D alignment was used to compare the newly identified viral RdRPs with known viral RdRPs, eukaryotic RdRPs, eukaryotic DdRPs and RTs to determine their degree of structural similarity (Figure 4D). The new RdRP supergroups from LucaProt contained at least three signature components of viral RdRP structures that resulted in significantly higher structural similarity to known RNA virus proteins (average structure similarity=3.0) than their cellular counterparts (average structure similarity=5.8).

**Figure 4.**
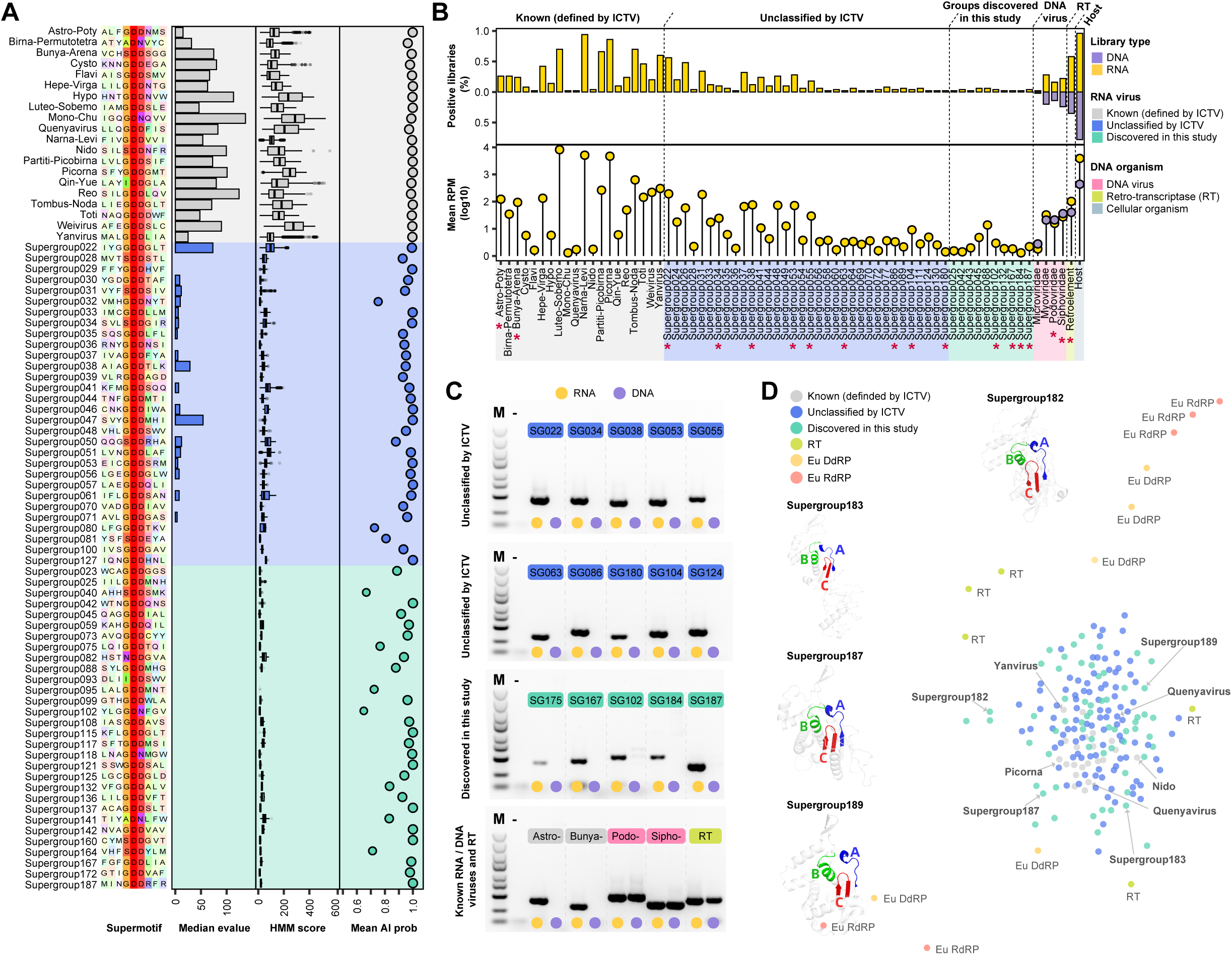
Evaluation of authenticity of RNA viral supergroups. (A) Distribution of BLAST median *e*-value, HMM score and mean AI modeling probabilities of RNA virus supergroups, with the conserved RdRP motif C of each supergroup shown on the left. For each group selected, the top 20 to 30 supergroups are shown by size for clarity. The known viral supergroups defined by the ICTV show high sensitivity for all three methods and are shaded in grey. The unclassified supergroups and new viruses discovered in this study show declining homology but a relatively stable AI probability. (B) The proportion of positive libraries and mean RPM (i.e., the number of mapped reads per million non-rRNA reads) of representative viral supergroups, DNA viruses, RT, and cell organisms in 50 samples collected in this study. DNA libraries are shown in purple and RNA libraries in yellow, while the different groups of RNA viruses and DNA organisms are shown in different colors, and red asterisks refer to those subsequently validated by RT-PCR. (C) RT-PCR results of the first pairs of validation primers for representative RdRP sequences from 17 RNA viral supergroups, capsid sequences from two DNA viral families (*Podoviridae* and *Siphoviridae*), and RT sequences. (D) Three-dimensional (3D) structure similarity analysis of representative RdRPs from 180 viral supergroups with Eu DdRPs, Eu RdRPs, and RT. Each point denotes a representative structure. The distance between different points represents structural similarity: the greater the distance, the lower the structural similarity. Four RdRP domain structures of the AI-specific supergroups are displayed with the A, B and C motifs highlighted. Representative RdRP domain structures from all supergroups are available at https://doi.org/10.6084/m9.figshare.26298802.v14. See also Figures S4 and S5.

### Genomic structures reveal modularity and flexibility within the RNA virosphere

We next analyzed the composition and structure of the putative RNA virus genomes identified in this study. The length of the RdRP-encoding genomes or genome segments differed markedly within and between viral supergroups, although most were centered around 2,131 nt (Figure 5). Notably, however, our data set also contained exceptionally long RNA virus genomes identified from soil that belonged to the Nido-like supergroup. One of these genomes was 47.3 kb, making it one of the longest RNA viruses identified to date^37^ (Figures 5C and S6; Table S6). Interestingly, an additional ORF was identified between the 5’ end of the genome of this virus and the ORF encoding RdRP, although the function of the encoded protein is unclear due to a lack of sequence similarity to known proteins. In addition to the RdRP, we characterized the other predicted proteins encoded by the newly identified virus genomes. While most had no homologs in existing databases, we identified some that were related to structural (e.g., coat, capsid, glycoprotein and envelope proteins) and non-structural (e.g., helicase, protease, methyltransferase, movement protein, immune or host-related regulatory proteins) proteins from known viruses (Figure 5D). Importantly, the presence of these additional virus proteins in newly identified supergroups provided further evidence that they were from *bona fide* RNA viruses. Furthermore, the presence of phage-related proteins (i.e., phage coat, phage mat-A and phage integrase) indicates that some of the viruses likely infect prokaryotic hosts, although further validation is required. In addition, the occurrence of these proteins was incongruent with virus phylogenetic history as inferred from RdRP sequences (Figure 5E), indicative of a modular-like configuration of RNA virus genomes.

**Figure 5.**
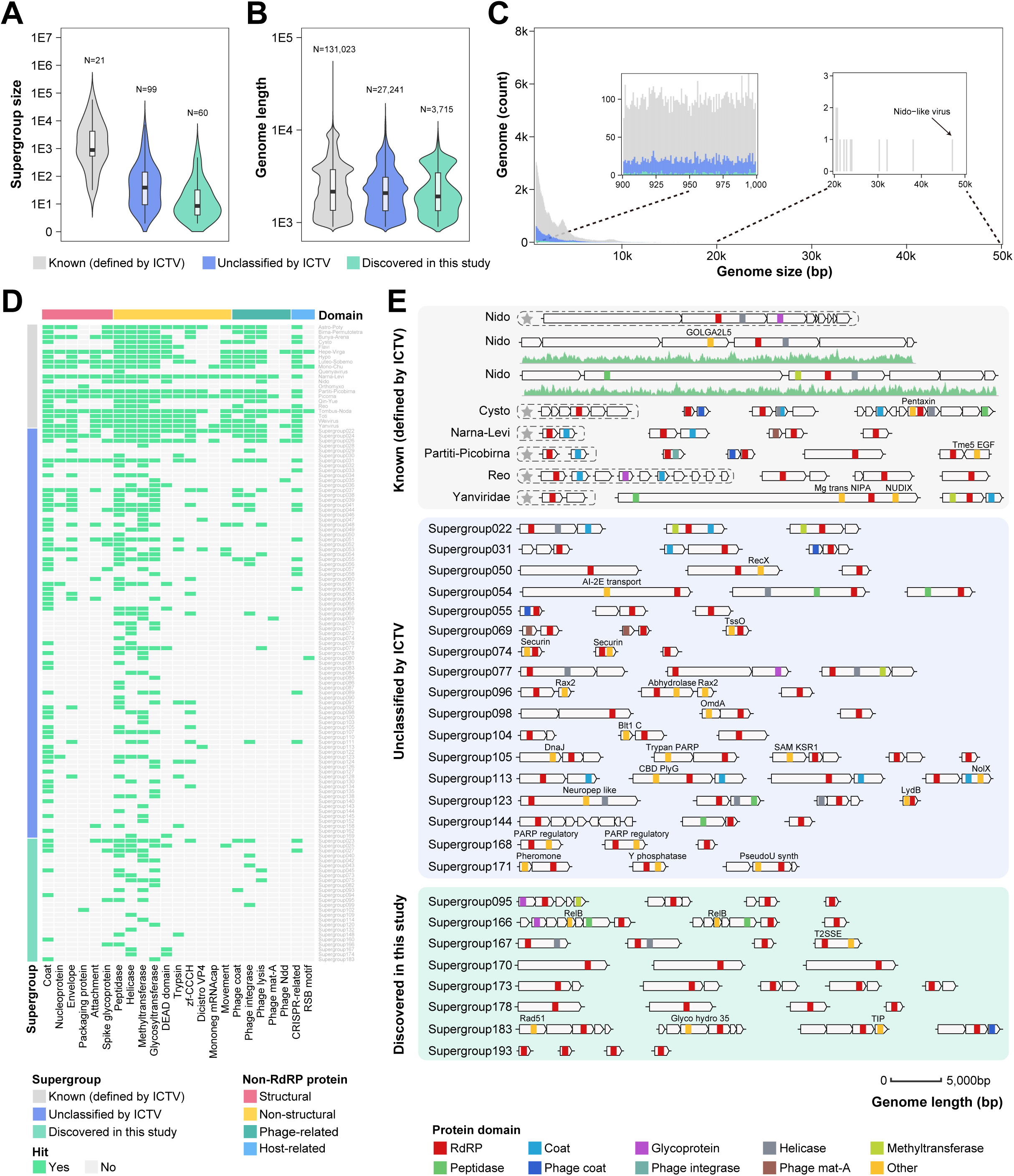
Genomic features of viral supergroups. (A) Size (i.e., the number of putative viral species) of all viral supergroups. The number of supergroups included in each group is displayed above the violin chart. Centre lines in the box plots represent the median bounds. (B) Genome length of viral species in all viral supergroups. The number of viral species included in each group is displayed above the violin chart. Centre lines in the box plots represent the median bounds. (C) Histogram of the genome size distribution of viral species from known, unclassified and new supergroups. (D) Distribution of annotated functional proteins in each viral supergroup. The supergroups with no annotated genes other than RdRP are not shown. (E) Genome structure of representatives from six known supergroups, 17 unclassified supergroups and eight new supergroups. Grey stars represent reference virus genomes of known supergroups. Domains not commonly found in RNA viruses are shown in yellow and are labeled above their corresponding positions. At the bottom, scale length in nucleotides. Abbreviations: GOLGA2L5: golgin subfamily A member 2-like protein 5; Pentaxin: pentaxin family; Tme5 EGF: thrombomodulin like fifth domain, EGF-like; Mg trans NIPA: magnesium transporter NIPA; NUDIX, nucleoside diphosphate-X hydrolase; RecX: RecX family; TssO: type VI secretion system, TssO; Securin: securin sister-chromatid separation inhibitor; Rax2: cortical protein marker for cell polarity; Abhydrolase: alpha/beta-hydrolase family; OmdA: bacteriocin-protection, YdeI or OmpD-Associated; Blt1 C: Get5 carboxyl domain; DnaJ: DnaJ domain; Trypan PARP: procyclic acidic repetitive protein (PARP); SAM KSR: kinase suppressor RAS 1; CBD PlyG: PlyG cell wall binding domain; LydB: LydA-holin antagonist; RelB: RelB antitoxin; T2SSE: type II/IV secretion system protein; PARP regulatory: poly A polymerase regulatory subunit; Pheromone: fungal mating-type pheromone; Y phosphatase2: tyrosine phosphatase family; PseudoU synth: RNA pseudouridylate synthase; Glyco hydro 35: glycosyl hydrolases family 35; TIP: tuftelin interacting protein. See also Figure S6.

### Expanded phylogenetic diversity of RNA viruses

The expansion of the RNA virosphere at the level of virus species – a 55.9-fold increase (251,846/4,502) from those defined by ICTV and a 1.4-fold increase (251,846/181,388) from all previously described RdRP sequences – is also evident in both the enlargement of known virus groups (e.g., phyla, orders, families) and the identification of entirely novel groups (Figure 6). Many of viruses discovered here formed clades that were distinct from the lineages observed in previously described virus supergroups (Figure 6). Interestingly, several groups previously represented by only a limited number of genomes – namely the Astro-Poty, Hypo, Yan and Cysto clades – experienced a major expansion to encompass larger viral clusters with greater phylogenetic diversity (Figure 6). Several newly identified supergroups also had high levels of phylogenetic diversity, including SG023 (1,232 virus species), SG025 (466 virus species), and SG027 (475 virus species), suggesting that more highly divergent RNA viruses will be discovered in environmental samples. Following our analysis, the supergroups with the greatest number of virus species here were the Narna-Levi (58,063 virus species), Picorna-Calici (19,970 virus species), and Tombus-Noda (15,520 virus species).

**Figure 6.**
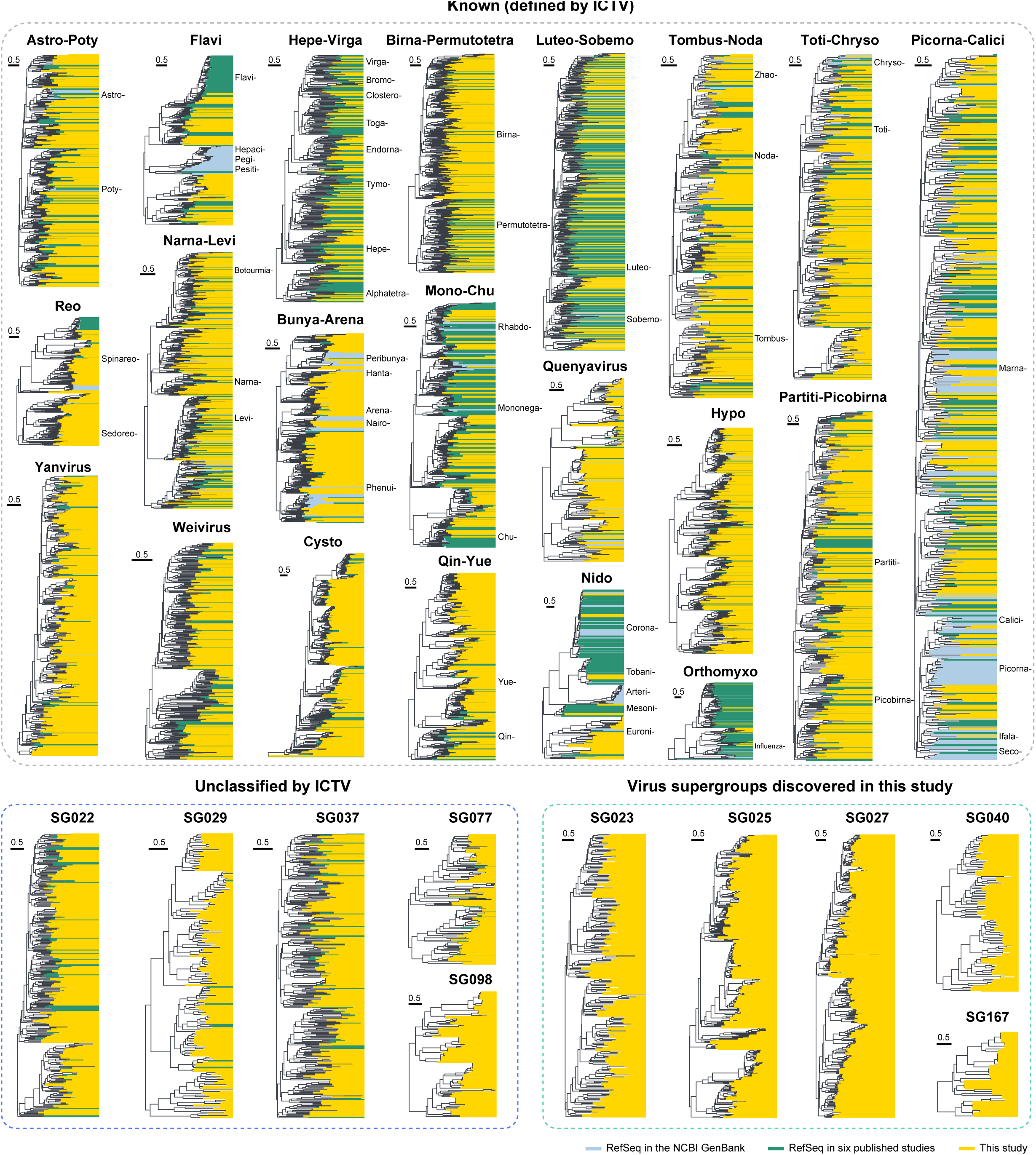
Phylogenetic diversity of 31 RNA viral supergroups. Each phylogenetic tree was estimated using a maximum likelihood method based on an amino acid alignment of the RdRP domain. Newly identified viruses are marked in yellow, those listed as ‘Viruses’ in the NCBI GenBank are marked in light blue, and viral RdRP sequences from six metagenomic studies^5–8,10,35^ are marked in green. The name of each supergroup is shown at the top of each phylogeny and the names of the families within each supergroup are shown to the right of the tree. All trees are midpoint-rooted for clarity only, and the scale bar indicates 0.5 amino acid substitutions per site. The tree files of all 180 supergroups are available at https://doi.org/10.6084/m9.figshare.26298802.v14.

### Ecological structure of the global RNA virome

Our analysis revealed the ubiquitous presence of RNA viruses across diverse ecosystem subtypes (32 categories) and in 1,612 locations globally. Despite repeated efforts to uncover the diversity of RNA viruses from such ecological samples,^5–8,10,35^ 33.3% of viral groups detected by LucaProt were not described previously (Figure 7A). Indeed, despite an overall deceleration, the rate of RNA virus discovery has not reached a plateau (Figure 7B), highlighting the vast untapped diversity of the global RNA virosphere. This is especially evident in soil environments, where there has been a notable increase in virus discovery.^10^

**Figure 7.**
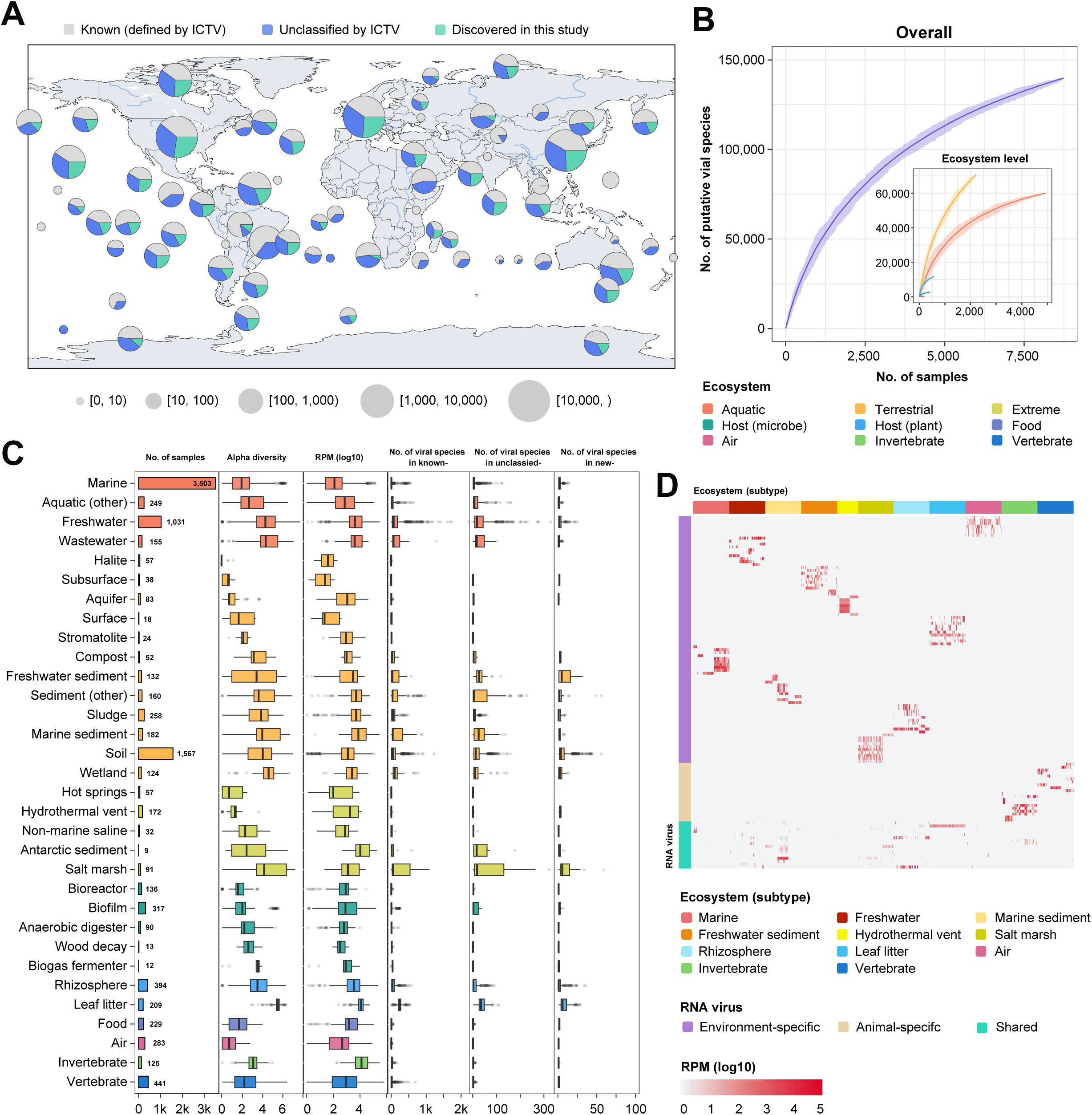
Ecological structure of the global RNA virome. (A) Global distribution of RNA viral species identified in this study. The Density-Based Spatial Clustering of Applications with Noise (NBSCAN) clustering algorithm^68^ was used to cluster the 1,612 different latitude and longitude points from positive samples. Points with a distance <800 km were aggregated, and the resulting 70 aggregated points were used to draw the map. Viral species from known RNA virus supergroups are shown in gray, from unclassified supergroups in blue, and from new supergroups in orange. Pie size reflects the number of viral species (log10). (B) Rarefaction curve of all RNA viral species. Inset, Rarefaction curve of RNA virus species at the ecosystem level with colors indicating different ecosystems. (C) Distribution of sample size (numbers in the bar chart), alpha diversity, RPM, virus species from known supergroups, virus species from unclassified supergroups and virus species from new supergroups at different ecosystem subtypes and colored by their ecosystem. “Extreme ecosystem” here refers to high salinity, high temperature, or low temperature environmental types. Only ecosystem subtypes with more than nine libraries (excluded poly-A sequencing) were retained for ecological comparison. The ecosystem subtypes on the y-axis are ordered from the highest to the lowest alpha diversity for each ecosystem. (D) Viral distributions in environmental and animal samples. The relative abundance of viruses in each library was calculated and normalized by the number of mapped reads per million no-rRNA reads (RPM). RNA virus species from 11 ecosystem subtypes are shown and placed into three groups, indicated by the colors on the heatmap. See also Figure S7.

To help identify any ecological patterns we compared alpha diversity (measured by the Shannon index) and abundance levels (measured by the number of reads per million total non-rRNA reads, i.e., RPM) of the RNA virome among diverse ecosystem subtypes (Figures 7C and 7D; Table S7). In general, average alpha diversity was highest in leaf litter, wetland, freshwater and wastewater environments, while virus abundance reached its peak in Antarctic sediment, marine sediment, and freshwater ecosystem subtypes, with average RPMs ranging from 18,424.6 to 46,685.5 (Figure 7C). In contrast, the lowest average diversity and abundance were in halite and subsurface environments (Figure 7C), which as expected due to their low biomass (i.e., host cells). For extreme ecological subtypes such as hot springs and hydrothermal vents, the associated RNA viruses were characterized by low diversity but moderate abundance (1,528.9 ∼ 3,726.9 average RPM) (Figure 7C). It is also noteworthy that the new viral supergroups established in this study were predominantly found in aquatic and sediment samples, with only a few occurrences in vertebrate and invertebrate animal samples (Figure 7C).

Our results further revealed the prevalence and abundance levels of particular viral groups across ecosystem subtypes (Figure 7D). Of note, the majority (85.9%) of the viruses discovered here only occurred in a single ecosystem subtype. Others, however, may be ecological generalists. For example, members of the Narna-Levi, Partiti-Picobirna and Picorna supergroups as well as Tombus-Noda supergroup were found in more than 32 ecosystem subtypes (Figure S7). Finally, we identified “marker” virus species, that exhibited high prevalence and abundance exclusively within specific ecosystem subtypes (Figure 7D), consistent with previous reports.^6,10^ Among these, *Partiti-Picobirna sp.*1991 and *Partiti-Picobirna sp.* 5447 were associated with hot springs, while *Tombus-Noda sp.* 2765 and *Supergroup 026 sp.* 2205 were associated with hydrothermal vents, suggesting a likely relationship with specialized hosts adapted to these environments.^38,39^ However, since the data sets analyzed here were generated by different laboratories employing distinct sample processing, library preparation and sequencing procedures, the comparisons of viral diversity and abundance among different ecosystem subtypes were necessarily subject to systemic biases.

## Discussion

The accurate identification of highly divergent RNA viruses remains a major challenge, impeding a comprehensive understanding of the genetic diversity in the RNA virosphere and hindering advancements in RNA virus evolution and ecology.^16,40^ Indeed, the conventional approach for RNA virus discovery has heavily relied on sequence similarity comparisons and the completeness of sequence databases.^40,41^ To address these issues, we developed a data-driven deep learning model (i.e., LucaProt) that outperforms conventional methods in accuracy, efficiency, and most importantly, the breadth of virus diversity detected. Importantly, LucaProt not only incorporated sequence data but also integrated structural information, which is crucial for accurate prediction of protein function, especially in the case of the RdRP.^42^ Without implementing the structural model, LucaProt had only 41.8% and 94.9% specificity and accuracy, respectively, on the test data set, and could only detect 44.5% of the predicted RdRP proteins. Hence, conservation of RdRP structure outweighs the significance of RdRP sequences in the identification of highly divergent RNA viruses. Collectively, we have established an AI framework paradigm for large-scale RNA virus discovery that can be readily extended to the accurate description of any biological “dark matter” once the training data set is prepared.

All the RNA viral sequences identified in this study were classified into clusters and supergroups, with the latter subsequently compared to viral classes and phyla as defined by the ICTV.^43,44^ Among the supergroups identified in this study, only 21 contained those from viral phyla/classes currently classified by ICTV, such that there was a 8.6-fold expansion of RNA virus diversity at the supergroup level compared to the latest ICTV report,^43^ and a 1.5-fold expansion of all RNA viruses described so far^5–8,10,35^ (Figure 1C). This expansion encompasses both existing viral supergroups as well as the discovery of 60 highly divergent supergroups that have largely been overlooked in previous RNA virus discovery projects (Figure 1D). The virus supergroups identified here were largely comparable to the existing classification system at the phylum (e.g., phylum *Lenarviricota* in the case of the Narna-Levi supergroup) or class (e.g., the *Stelpaviricetes, Alsuviricetes, Flasuviricetes* classes for the Astro-Poty, Hepe-Virga, Flavi supergroups) levels, highlighting the extent of the phylogenetic diversity identified here.

Despite the large expansion in RNA virus diversity documented here, major gaps remain in our understanding of the evolution and ecology of the newly discovered viruses. In particular, the hosts for most of the viruses identified remain unknown. As the majority of current known RNA viruses infect eukaryotes,^45,46^ and microbial eukaryotes exist in great abundance and diversity in natural environments,^47,48^ it is possible that the viral clades and supergroups identified here were largely associated with diverse microbial eukaryotic hosts. However, it is also likely that a substantial proportion of the novel viruses discovered are associated with bacterial (and perhaps archaeal) hosts.^49–51^ Indeed, mounting evidence^10,52^ strongly supports the notion that more groups of RNA viruses than currently documented are associated with bacteria. Indeed, the presence of RNA bacteriophage in multiple RNA viral supergroups underlines the evolutionary connection between RNA viruses from bacterial and eukaryotic hosts. If viewed through the lens of virus-host co-divergence,^1,2,53^ such a link suggests that the evolutionary history of RNA viruses is at least as long, if not longer, than that of the cellular organisms.

### Limitations of the study

Our study has several limitations. First, classifying viruses on a deep evolutionary scale remains a complex and challenging task due to the very high levels of genetic divergence exhibited by these viruses. This divergence is so pronounced that neither sequence nor structural homology can adequately uncover their true evolutionary histories. Second, while we successfully confirmed the RNA nature of some virus supergroups through RNA and DNA sequencing of the same sample, we could not apply this method to other viral supergroups. This limitation arose because these supergroups were identified using data from the SRA database that lack matching DNA sequencing data. Finally, the viral genomes we identified only contain segments associated with the RdRP. For segmented viruses, this means we only uncovered partial genomes, with the greater sequence divergence in non-RdRP segments hindering their identification. Future efforts using more generalized models could potentially identify all viral segments. AI models could be expanded to detect viral functional proteins beyond RdRP, enabling the identification of segments encoding highly divergent viral proteins. Alternatively, as all viral genome segments must have broadly matching levels of abundance (i.e. co-occur) as the RdRP, the detection of sequences at similar abundance to the RdRP could be used to identify additional segments, particularly if this pattern of co-occurrence is found in multiple libraries.

## ACKNOWLEDGMENTS

This work was supported by the National Natural Science Foundation of China (82341118, 32270160), the Shenzhen Science and Technology Program (KQTD20200820145822023 and JCYJ20210324124414040), the Natural Science Foundation of Guangdong Province (2022A1515011854), the Guangdong Province “Pearl River Talent Plan” Innovation and Entrepreneurship Team Project (2019ZT08Y464), the Hong Kong Innovation and Technology Fund (ITF) (MRP/071/20X), and the Health and Medical Research Fund (COVID190206). E.C.H. is funded by a National Health and Medical Research Council (Australia) Investigator grant (GNT2017197) and by AIR@InnoHK administered by the Innovation and Technology Commission, Hong Kong Special Administrative Region, China.

We thank the Computing and Storage teams of Alibaba Cloud Computing Co., Ltd. and Zhejiang laboratory for their contribution of 15 machines with 128 CPUs and 1T RAM of Elastic High-Performance Computing (EHPC), 64 Nvidia A100 Graphics Processing Units (GPUs), and 500TB of Network Attached Storage (NAS) resources.

## AUTHOR CONTRIBUTIONS

Conceptualization, X.H., Y.H., E.C.H., Z.-R.L. and M.S.; Methodology, X.H., Y.H., J.-S.E., J.L., Z.-R.L. and M.S.; Investigation, X.H., Y.H., P.F., S.-Q.M., Z.X. and Q.-Y.G.; Writing – Original Draft, X.H., Y.H., E.-C.H. and M.S.; Writing – Review and Editing, All authors. Funding Acquisition, F.-M.H., Y.-L.S., D.-Y.G., Z.-R.L. and M.S.; Resources (sampling), X.H., S.-Q.M., W.-W.C., J.-H.T., G.-Y.X., S.-J.L., Y.-Y.X., Y.-L.Z., F.-M.H., Y.-F.P., Z.-H.Y. and C.H.; Resources (computational), S.Z., Z.-Y.Z. and Z.-R.L.; Supervision, Z.R.L. and M.S.

## Supplemental figure titles and legends

**Figure S1.**
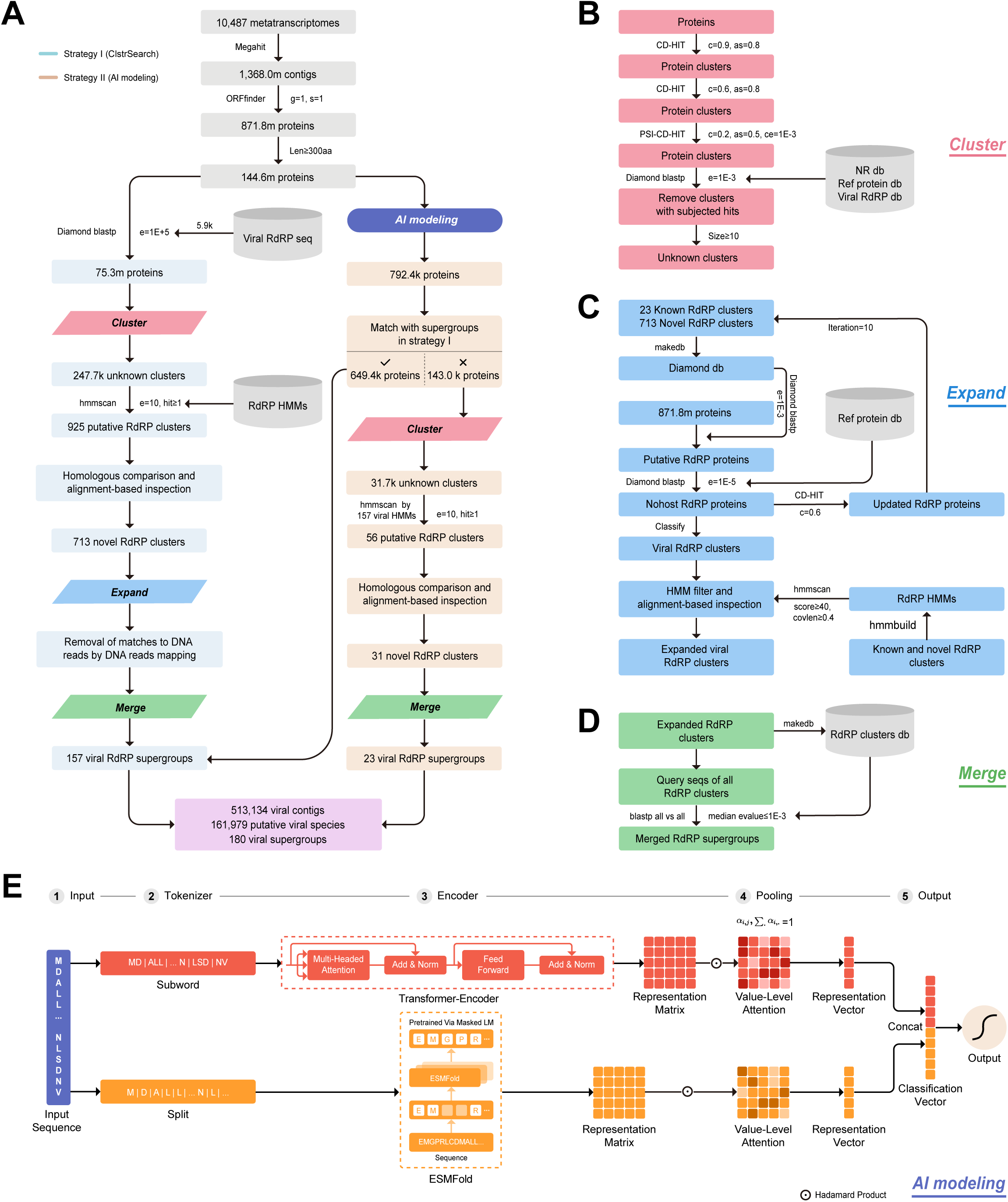
Detailed RNA virus discovery pipeline, related to Figure 1. (A) Schematic diagram of the similarity-based discovery (i.e., ClstrSearch) and RdRP AI modeling (i.e., LucaProt) approaches. (B) Protein clustering process. Only clusters with more than ten members are retained for viral cluster discovery. (C) Ten iterations of RdRP expansion by recruiting newly detected RdRPs. (D) RdRP clusters are merged into RdRP supergroups using BLAST median *e*-values. (E) RdRP identification by LucaProt that includes five modules: Input, Tokenizer, Encoder, Pooling, and Output.

**Figure S2.**
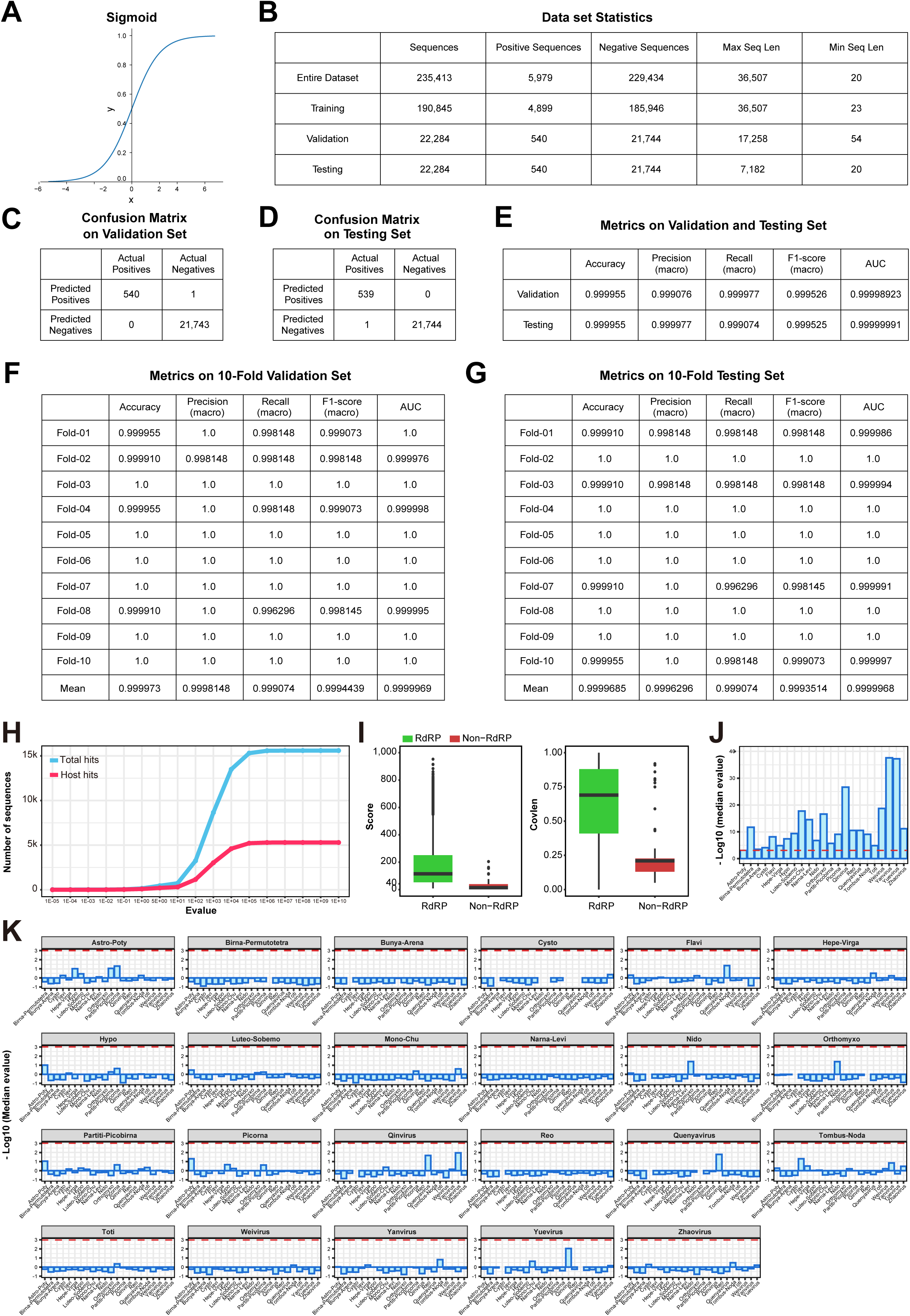
Benchmarking of LucaProt for the RdRP modeling and threshold used in ClstrSearch, related to Figure 1. (A) The Sigmoid function of the LucaProt. The Sigmoid function (i.e., the Logistic function) is used for the output layer of a binary classification model and for mapping a real number to the probability of 0 ∼ 1. (B) Statistics of the data set used for LucaProt building, including the entire data set, training set, validation set, and testing set. (C) The confusion matrix on the validation data set. (D) The confusion matrix on the testing data set. (E) The metrics for the validation and testing data sets. (F) and (G) The metrics on the validation and testing sets of 10-fold cross-validation for LucaProt. The positive and negative samples were randomly shuffled and divided into ten parts, with each part containing 540 positive and 21,744 negative samples. In the 10-fold cross-validation, two parts were selected rotationally to serve as the validation and testing sets, respectively, while the remaining samples were combined to form the training set. (G) Number of hits with Diamond blastp v0.9.25.126 using different *e*-values at the test stage. A total of 15,000 sequences were randomly sampled for the test, and host hits were recognized by comparison against the NCBI non-redundant (nr) protein database (version 2023.01.09) with an *e*-value threshold of 1*E*-3. (I) Benchmarking of HMMscan bitscore and aligned fraction using the RdRP and non-RdRP data sets (including RT, Eu DdRP and Ed RdRP derived from the NCBI GenBank database). (A) (J) BLAST Median *e*-value within the same known RdRP cluster. (K) BLAST Median *e*-value between pairwise comparisons of known RdRP clusters, with a 1*E*-3 cut-off used for cluster merging.

**Figure S3.**
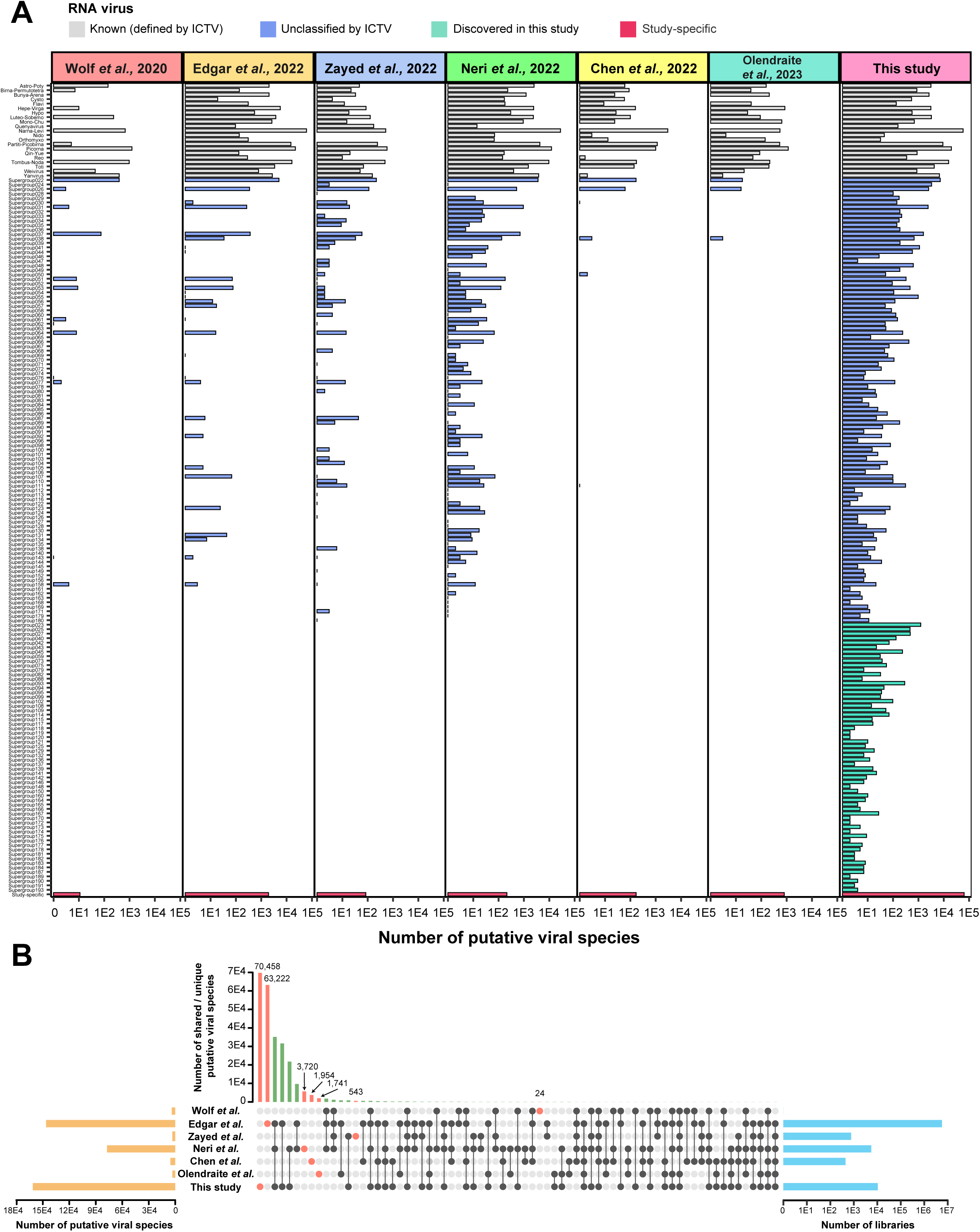
Comparison of RNA virus discovery results between six previous studies and the current study, related to Figure 1. (A) The distribution of putative viral species of seven studies at the supergroup level and the study-specific level. (B) Upset plot showing the number of viral species found in each study and those that are shared/unique between and among seven studies. The red bar in the top histogram represents unique viral species identified in each study, while the green bar indicates viral species that are shared between two or more studies. Rightmost histogram shows the number of metatranscriptomes analyzed in each study. Of note, Olendraite *et al*. sourced viral RdRPs from the NCBI Transcriptome Shotgun Assembly (TSA) database, such that the exact number of metatranscriptomes analyzed is unclear.

**Figure S4.**
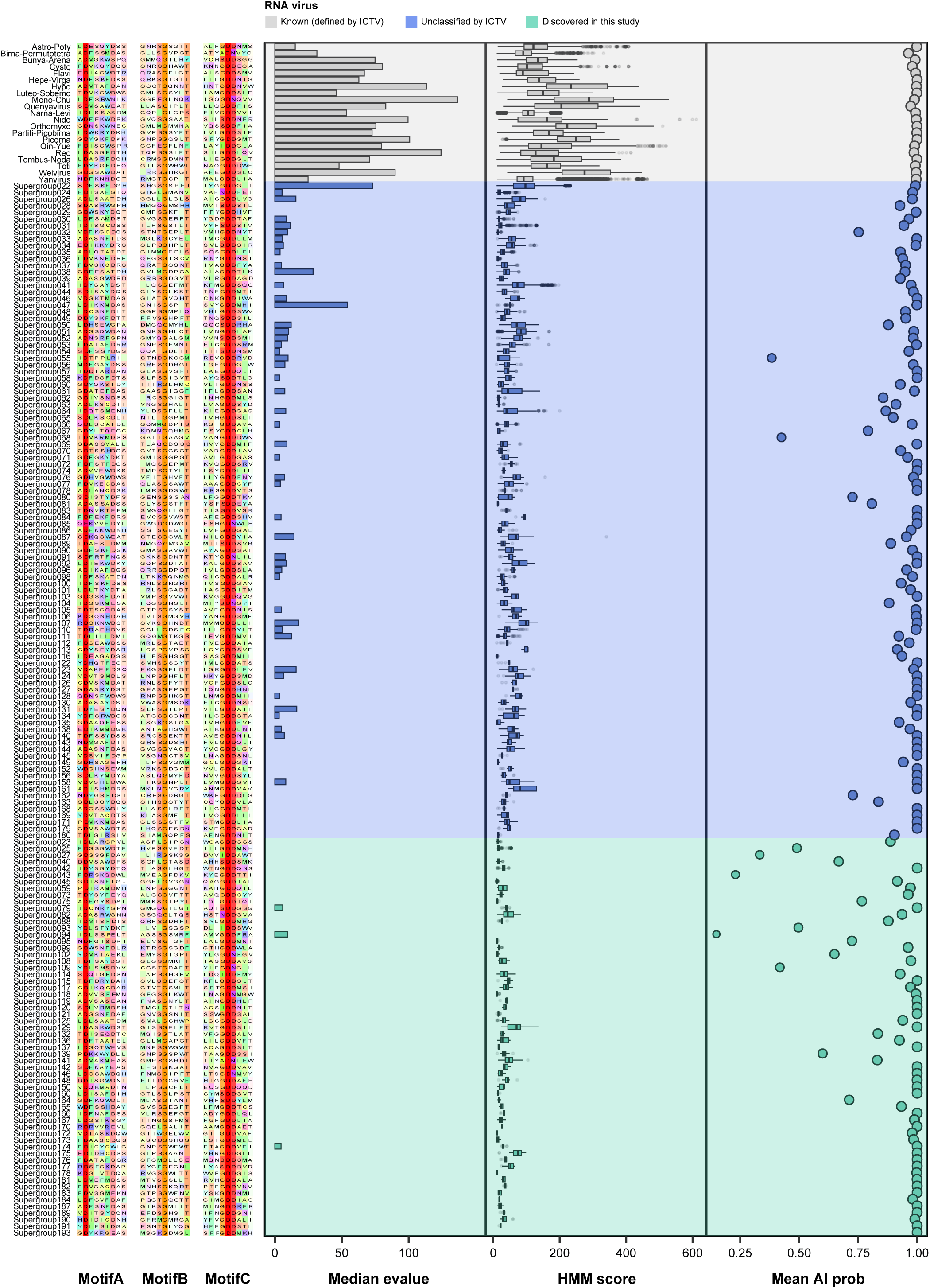
The three highly conserved RdRP sequence motifs (A, B and C) and the distribution of BLAST median e-value, HMM score and mean AI modeling probabilities of the RNA virus supergroups, related to Figure 4. RNA viral supergroups were divided into three groups by similarity to reference and other published data sets, with the conserved RdRP A, B and C motifs of each supergroup shown on the left.

**Figure S5.**
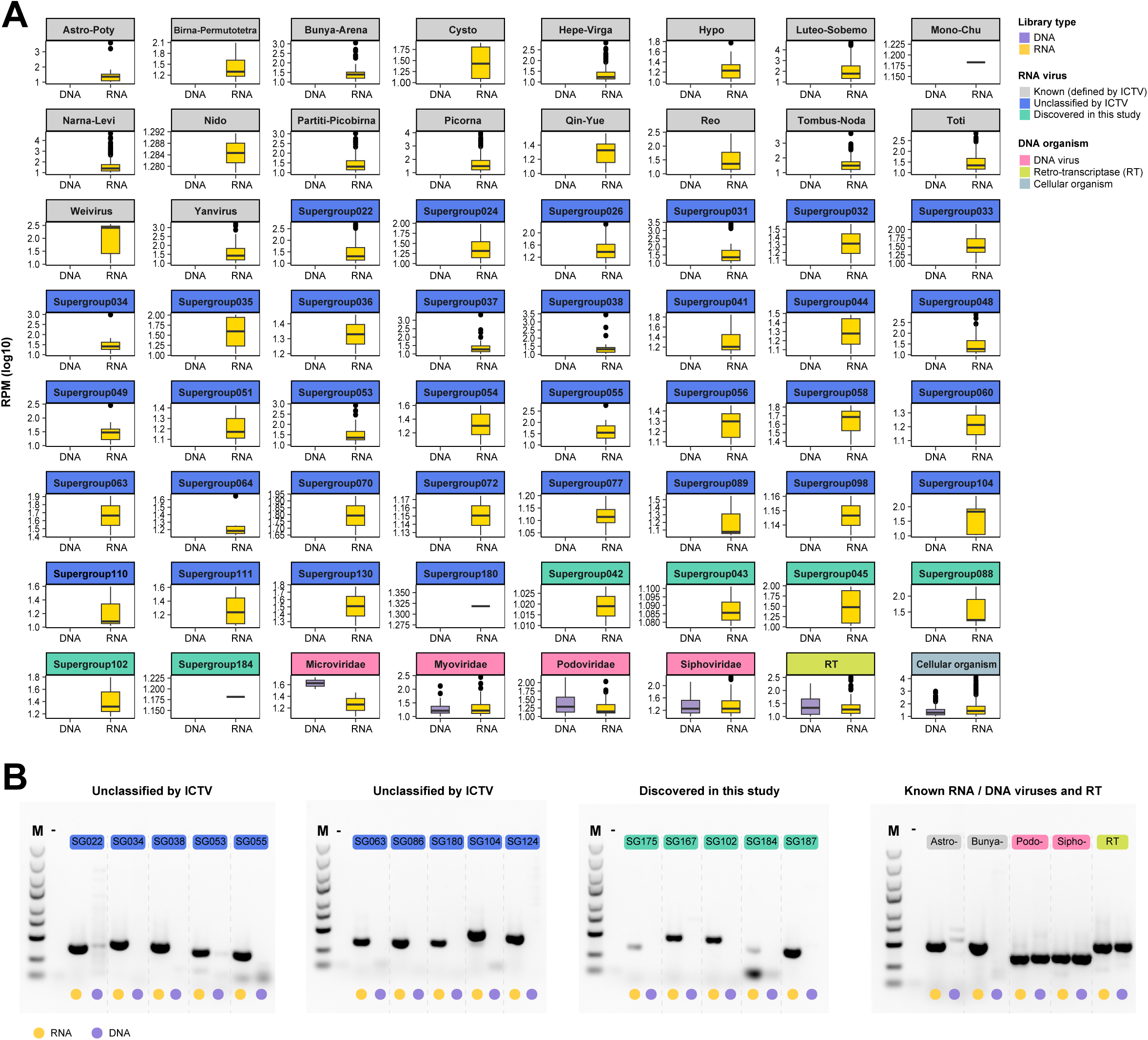
Expression difference of RNA viruses and DNA organisms in our newly sequenced data, related to Figure 4. (A) Abundance comparisons for 58 RNA viral supergroups, four DNA virus families, RT and cell organisms at DNA and RNA libraries. (B) RT-PCR results of second pairs of validation primers for representative RdRP sequences from 17 RNA viral supergroups, capsid sequences from two DNA virus families (*Podoviridae* and *Siphoviridae*), and RT sequences.

**Figure S6.**
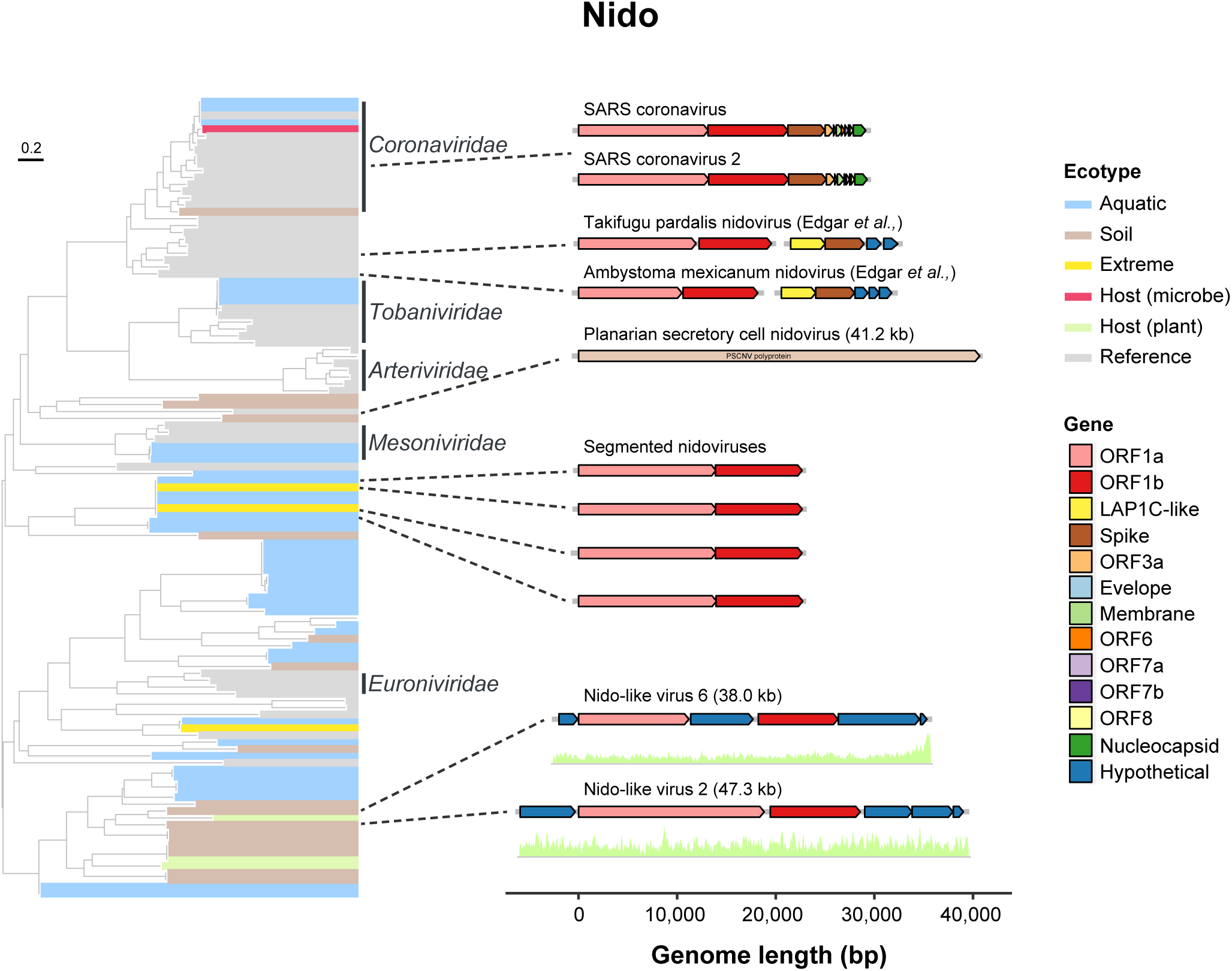
Phylogenetic tree of the Nido-like supergroup and the genome structure of representative viruses, related to Figure 5. The tree was estimated using a maximum likelihood method based on the conserved RdRP domain. The reference sequences reported previously are shaded grey, while the viruses newly identified here are shaded by different colors according to ecotype. The names of viral families are shown on right of the tree. The tree was midpoint-rooted for clarity only, and the scale bar of tree indicates 0.2 amino acid substitutions per site. The genome structures of representative viruses are shown on right of the tree. The scale bat at the bottom indicates the length in nucleotides.

**Figure S7.**
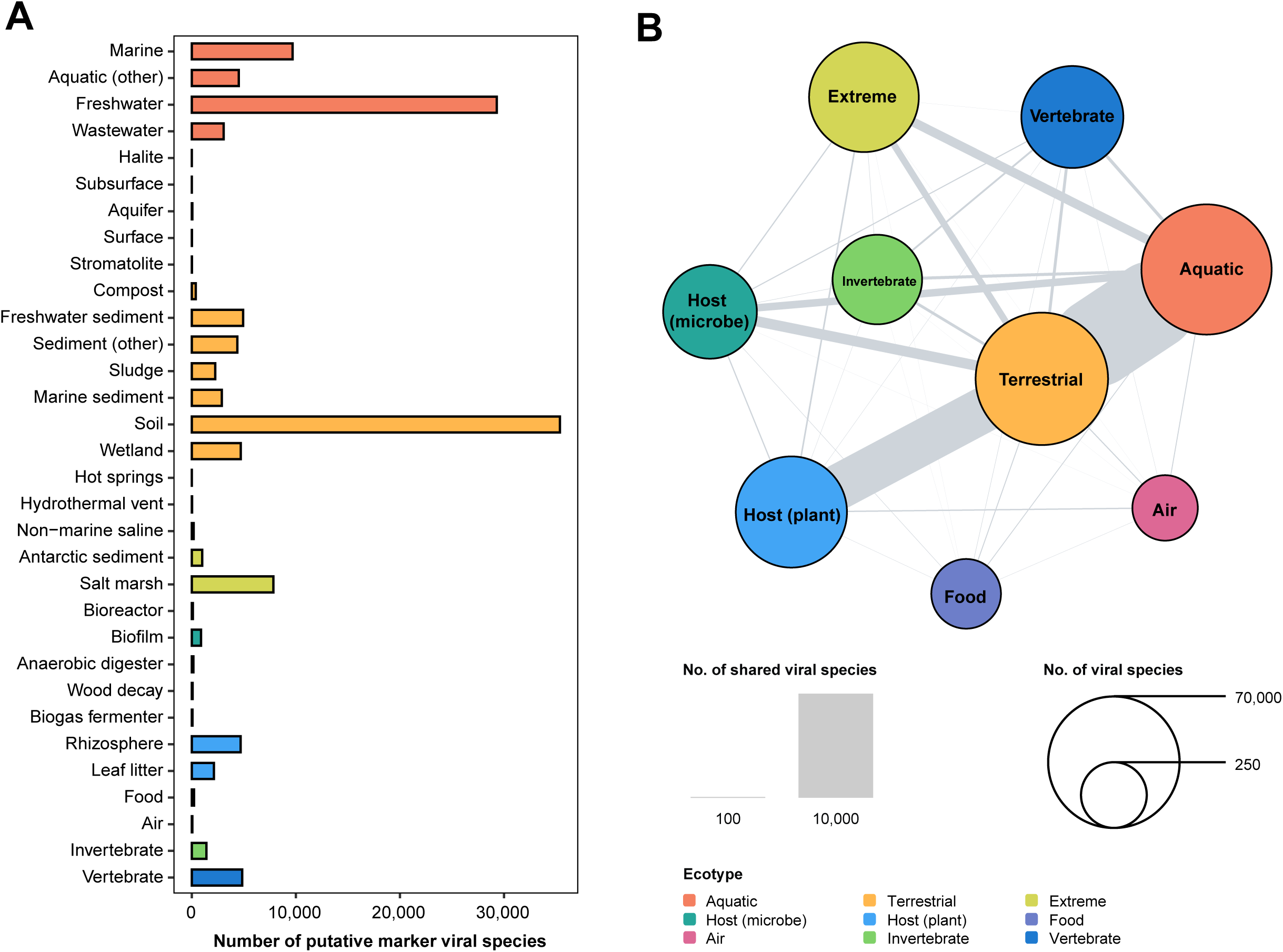
Specificity and shareability of RNA viruses, related to Figure 7. (A) Number of specific putative viral species (i.e., putative “marker” viral species) in each ecosystem subtype. (B) Association between RNA viruses and different environmental ecosystems. The size of the colored circles indicates the number of putative viral species identified by each ecosystem type, while the thickness of the line indicates the number of viral species shared by each ecosystem.

## STAR*METHODS

Detailed methods are provided in the online version of this paper and include the following:

- KEY RESOURCES TABLE
- RESOURCE AVAILABILITY

- Lead Contact
- Materials availability
- Data and code availability
- METHOD DETAILS

- Samples and data sets
- Identification of RNA viruses based on deep learning
- Identification of RNA viruses based on homologous clustered proteins
- Benchmarking LucaProt and comparisons to Diamond, HMMscan, HH-suite and PalmScan
- Virus verification
- Structural prediction and comparisons between viral RdRPs and homologous proteins
- Annotation and characterization of virus genomes
- Analyses of virome diversity, evolution and ecology
- QUANTIFICATION AND STATISTICAL ANALYSIS

## SUPPLEMENTAL INFORMATION

Supplemental information can be found online at XXXXXXXXXX

## STAR*METHODS

### KEY RESOURCES TABLE

| REAGENT or RESOURCE | SOURCE | IDENTIFIER |
| --- | --- | --- |
| RNeasy®PowerSoil® Total RNA Kit | QIAGEN | Cat. #12866-25 |
| RNeasy® PowerSoil® DNA Elution Kit | QIAGEN | Cat. #12867-25 |
| NEB Next Ultra RNA Library Prep Kit | New England Biolabs | Cat. #E7770 |
| NEB Next Ultra DNA Library Prep Kit | New England Biolabs | Cat. #E7370L |
| **Biological samples** |  |  |
| Samples are described in Table S2 | This paper | N/A |
| **Deposited data** |  |  |
| Raw sequencing reads generated in this study | This paper | NCBI-SRA BioProject: PRJNA956286 (RNA-seq) and PRJNA956287 (DNA-seq) |
| All assembled contigs used in this study | This paper | https://ftp.cngb.org/pub/CNSA/data4/CNP0005901/CNS1149693/CNA0141908/ |
| Representative viral genome sequences | This paper | https://db.cngb.org/search/project/CNP0005901/ |
| All original data and code produced in this work | This paper | https://doi.org/10.6084/m9.figshare.26298802.v14 |
| Original code produced in this study | This paper | https://github.com/alibaba/LucaProt |
| **Software and algorithms** |  |  |
| MEGAHIT v1.2.8 | Li et al.<sup>54</sup> | https://github.com/voutcn/megahit |
| ORFfinder v0.4.3 | NCBI Team | https://ftp.ncbi.nlm.nih.gov/genomes/TOOLS/ORFfinder/linux-i64/ |
| ESMFold | Lin et al.<sup>56</sup> | https://github.com/facebookresearch/esm |
| Subword-NMT v0.3.8 |  | https://github.com/rsennrich/subword-nmt |
| Diamond v0.9.25.26 | Buchfink et al.<sup>58</sup> | https://github.com/bbuchfink/diamond |
| CD-HIT v4.8.1 |  | https://github.com/weizhongli/cdhit |
| HMMER | Potter et al.<sup>59</sup> | http://hmmer.org/ |
| Mafft v7.475 | Katoh et al.<sup>60</sup> | https://mafft.cbrc.jp/alignment/software/ |
| Girvan-Newman algorithm | Girvan et al.<sup>62</sup> | https://memgraph.github.io/networkx-guide/algorithms/community-detection/girvan-newman/ |
| iGraph package v1.3.5 | Csárdi et al.<sup>63</sup> | https://igraph.org |
| HH-suite v3.3.0 | Steinegger et al.<sup>64</sup> | https://github.com/soedinglab/hh-suite |
| PalmScan | Babaian et al.<sup>61</sup> | https://github.com/rcedgar/palmscan |
| Bowtie2 v2.4.2 | Langmead et al.<sup>65</sup> | https://bowtie-bio.sourceforge.net/bowtie2/index.shtml |
| AlphaFold2 v2.3 | Jumper et al.<sup>20</sup> | https://alphafold.com/ |
| PyMol v2.5.4 |  | http://www.pymol.org/pymol |
| NetworkX |  | https://networkx.org/ |
| PhyML v3.1 | Guindon et al.<sup>67</sup> | http://www.atgc-montpellier.fr/phyml/ |

### RESOURCE AVAILABILITY

#### Lead Contact

Further information and requests for resources and additional data should be directed to and will be fulfilled by the Lead Contact, Mang Shi.

#### Materials availability

This study did not generate new unique reagents or specific biological material. The materials used and generated in this study are available from the Lead Contact. As a computational project, the input, results, output, and code of this study is publicly available and are described below in the ‘‘data and code availability’’ section.

#### Data and code availability

The raw sequence reads newly generated in this study are available at the NCBI Sequence Read Archive (SRA) database under BioProject accessions PRJNA956286 and PRJNA956287 (Table S2). All assembled contigs from this study are available in the China National GeneBank DataBase (CNGBdb) under the accession CNP0005901. The representative viral genome sequences generated are available in the CNGBdb under the accession CNP0005901 (https://db.cngb.org/search/project/CNP0005901/). Additionally, the related result data sets and LucaProt-related resources are publicly available at https://doi.org/10.6084/m9.figshare.26298802.v14, including: (i) the result data sets from this study, including viral contigs, viral RdRP sequences, viral RdRP HMM profiles and phylogenetic tree files; (ii) the LucaProt code, the data sets for model building, and the trained model of LucaProt; (iii) the inference and prediction code using the trained model for unknown sequences, benchmarks with other methods, and (iv) additional resources.

The original source code for ClstrSearch and LucaProt are stored at the GitHub repository (ClstrSearch: https://github.com/alibaba/LucaProt/tree/master/ClstrSearch, LucaProt: https://github.com/alibaba/LucaProt). We also provide the Python dependency environment installation file, installation commands, and the running command of the trained LucaProt model for inference or prediction, which can be found in the same GitHub repository. These models are compatible with Linux, Mac OS, and Windows systems, supporting both CPU and GPU configurations for inference tasks.

### METHOD DETAILS

#### Samples and data sets

This study comprised RNA virus discovery through the metatranscriptomic analysis of 10,487 samples. The majority of these samples (n=10,437) were mined from the NCBI Sequence Read Archive (SRA) database (https://www.ncbi.nlm.nih.gov/sra) between January 16 - August 14, 2020. We targeted samples collected from a wide range of environmental types globally (Figure 2), including: aquatic (such as marine, riverine and lake water), soil (such as sediment, sludge and wetland), host-related (such as biofilm, wood decay, and rhizosphere), and extreme environmental samples (such as hydrothermal vent, hypersaline lake and salt marsh), that were subject to high quality metatranscriptomic sequencing to ensure the generation of ≥50 Mb total RNA Q20 sequencing data. In addition, 50 data sets were generated in this study (see below), all of which were subject to high-quality short-read sequencing utilizing Illumina sequencing platforms. The raw sequencing data output ranged from 35.1 to 204.1 Gbp, and no enrichment for microbial organisms was performed during sample processing or library preparations. For highly abundant environmental types, such as “soil” and “marine”, representative samples were selected to include as many projects (i.e., independent studies), geographic locations and ecological niches as possible.

In addition to data mined from the SRA database, we generated 50 metatranscriptomic data sets from Antarctica and China. The sample types included marine (n=5), freshwater (n=12), soil (n=19), and sediment (n=14), of which nine sediment samples were collected at the Ross Sea station in Antarctica between January and February 2022, with the others from Zhejiang, Guangdong, Hubei and Heilongjiang provinces of China collected between August and October 2022. For each of these samples, DNA and RNA were simultaneously extracted: the soil and sediment samples were extracted using the RNeasy® PowerSoil® Total RNA Kit and RNeasy® PowerSoil® DNA Elution Kit (QIAGEN, Germany), while the marine and freshwater samples were extracted using the DNeasy® PowerWater® Kit and RNeasy® PowerWater® Kit (QIAGEN, Germany). The extracted nucleic acid was then subject to library construction using the NEB Next Ultra RNA Library Prep Kit and NEB Next Ultra DNA Library Prep Kit (New England Biolabs, China) for RNA and DNA samples, respectively. Paired-end (150 bp) sequencing of these libraries was performed using the Illumina NovaSeq 6000 platform (Illumina, USA).

For all 10,487 data sets used this study, sequence reads were *de novo* assembled into contigs using MEGAHIT v1.2.8^54^ with default parameters, without implementing any quality control procedures to optimize time efficiency. Potential encoded proteins were predicted from contigs using ORFfinder v0.4.3 (https://ftp.ncbi.nlm.nih.gov/genomes/TOOLS/ORFfinder/linux-i64/; parameters, -g 1, -s 2).

### Identification of RNA viruses based on deep learning

We developed a new deep learning, transformer-based model, termed “Deep Sequential and Structural Information Fusion Network for Protein Function Prediction” (i.e., LucaProt), that effectively integrated protein sequence composition and structure information to identify viral RdRPs. The model contains two channels. The first channel employs a Transformer-Encoder for feature extraction from the raw sequence data. We used the Byte-Pair Encoding (BPE) algorithm^55^ for tokenization, enabling the model to handle longer sequences and identify meaningful units (such as words, rather than individual residues) within sequences associated with RdRPs. The BPE algorithm creates a word-level vocabulary from a sequence corpus and tokenizes the input sequence accordingly, prior to feature extraction via the Transformer-Encoder. The second channel leverages the protein language model ESM2-3B^56^ to capture residue-level features, utilizing the model’s self-supervised learning capabilities to decode contextual structural information. The model concludes with two Value-Level Attention Pooling layers that down-sample each feature matrix into a vector. These two vectors are then concatenated for the final classification task. The model includes five modules: Input, Tokenizer, Encoder, Pooling, and Output (Figure S1E).

#### Input Layer

This layer accepts amino acid sequences as the model’s input.

#### Tokenizer Layer

This layer processes raw sequence data into a digestible format for feature extraction. This module comprises two components. First, a corpus comprising 5,979 positive sequences (i.e., viral RdRPs) and 150,000 sequences sampling from negative samples (i.e., protein sequences that are not viral RdRPs) was constructed. The BPE algorithm^55^ from the Subword-NMT tool (https://github.com/rsennrich/subword-nmt) was utilized to create a vocabulary of about 20,000 frequent subsequences or “words”. These words were derived by treating co-occurring amino acids in the sequences as single entities. Additionally, each protein sequence was broken down into individual amino acids for subsequent extraction of protein structural information using a large protein language model.

#### Encoder Layer

This layer transforms the data into two representations by different tokenization strategies (as illustrated in the former layer): a sequence representation matrix with word-level tokenization and a structural representation matrix with character-level tokenization. For sequence processing, an advanced Transformer-Encoder model was utilized to generate the sequence representation matrix. In terms of structural processing, the intermediate matrices produced by the structural prediction model ESMFold (ESM2-3B)^56^ were used as the structural representation matrix. This methodology addresses the limitation posed by the scarcity of experimentally observed 3D structures and obviates the need for additional encoding steps, thereby guaranteeing computational efficiency.

#### Pooling Layer

This layer reduces dimension and selects features for efficient classification. It employs a Value-Level Attention Pooling (VLAP) approach^57^ to convert the sequence and structural matrices into two vectors.

#### Output Layer

This layer transforms the pooled vectors into the probability value indicative of a positive sample. The vectors from the pooling layer are concatenated and passed through a fully connected layer. A sigmoid function outputs probability values ranging from 0.0 to 1.0, with a threshold of 0.5 to classify sequences as viral RdRP or not (Figure S2).

#### Model Building

We constructed a data set of 235,413 samples for model building. This included 5,979 positive samples of known viral RdRPs (i.e., the well-curated RdRP database described below) and 229,434 randomly selected negative samples of confirmed protein sequences that were not viral RdRPs (as the positive samples account for a very small portion of the total data, we constructed the training data set using the conventional 1:40 ratio of positive to negative data). The representative negative samples contained proteins from eukaryotic RNA-dependent RNA polymerase (Eu RdRP, n=2,233), eukaryotic DNA-dependent RNA polymerase (Eu DdRP, n=1,184), reverse transcriptase (RT, n=48,490), proteins obtained from DNA viruses (n=1,533), non-RdRP proteins obtained from RNA viruses (n=1,574), as well as an array of cellular proteins from different functional categories (n=174,420). We randomly divided the data set into training, validation, and testing sets with a ratio of 8.5:1:1, which were used for model fitting, model finalization (based on the best F1-score training iteration), and performance reporting (including accuracy, precision, recall, F1-score, and Area under the ROC Curve (AUC)), respectively (Figure S2).

We further validated the performance and robustness of LucaProt model by implementing a 10-fold cross-validation (Figure S2). During this process, both positive and negative samples were randomly shuffled and divided into ten equal parts, with each part comprising 540 positive and 21,744 negative samples. For the cross-validation, two parts were sequentially selected as the validation and testing sets, respectively, while the remaining samples were merged to form the training set.

LucaProt successfully identified 792,436 putative RdRP signatures from a data set of 144.6 million proteins derived from the 10,487 metatranscriptomes. These results were first compared with the RdRPs identified based on ClstrSearch (see below). RdRPs that were identified by deep learning algorithms (i.e., LucaProt) were either incorporated into RNA virus supergroups using the Diamond blastp program v0.9.25.126^58^ with an *e*-value threshold of 1*E*-3, or, if they remained unclassified, were subjected to clustering, merging, and manual alignment inspection. The presence of key viral RdRP motifs – motif A [DxxxxD], motif B [(S/T)Gxxx(T/G)xxxN], and motif C [(S/G/N/A)D(D/N)] – were examined for all supergroups through multiple sequence alignments.

### Identification of RNA viruses based on homologous clustered proteins

Another strategy for the identification of RNA viruses at the cluster level involved the utilization of homologous clustered proteins (Figure S1A). Accordingly, a total of 871.8 million amino acid sequences predicted by ORFfinder (see Samples and data sets) were compared against a well-curated RdRP database (n=5,979) that contained only those derived from reference RNA virus genomes downloaded from the NCBI GenBank database and their close relatives in vertebrate and invertebrate hosts.^1,2^ The comparisons were performed using the Diamond blastp program v0.9.25.126,^58^ with the *e*-value threshold set at 1*E*+5 to identify more divergent RdRP sequences (Figures S1A and S2H). This process resulted in 75.3 million hits that were further subjected to similarity-based and multi-step clustering (three iterations with 90%, 60%, and 20% amino acid identity, respectively) using CD-HIT v4.8.1 (https://github.com/weizhongli/cdhit); this process resulted in 3,805,584 clusters. False positives and hits to known RdRP sequences were removed by comparison against the NCBI non-redundant (nr) protein database (version 2023.01.09), the NCBI RefSeq protein database (version 2023.03.26) and the virus RdRP database using the Diamond blastp program v0.9.25.126^58^ with an *e*-value threshold of 1*E*-3 (Figure S1B). The remaining unknown protein clusters were subject to viral RdRP domain search using hidden Markov models (HMMs) built from a manually reviewed profile of known RdRP clusters using HMMscan v3.3.2 (*e*=10, hits≥1).^59^ The RdRP sequences of each known viral group were aligned using Mafft v7.475,^60^ employing the L-INS-I algorithm. HMM profiles were then built from each MSA using HMMbuild v3.3.2 with standard parameters.^59^ Clusters that contained more than one HMMscan hit were subsequently aligned and inspected for the presence of well-conserved RdRP core sequences referred to as the “palmprint”, delineated by the presence of the three motifs (A, B and C) that collectively constitute the catalytic core within the RdRP structure.^61^ As a result of our rigorous screening and checking steps, a total of 713 potential RdRP clusters were retained.

To further expand the RdRP collection based on the viruses newly discovered here, we updated the RdRP protein database with the 713 RdRP clusters newly identified here and used it to detect additional RdRP sequences from the original 144.6 million amino acid sequences using Diamond blastp v0.9.25.126^58^ with an *e*-value threshold of 1*E*-3. The newly detected RdRPs were again incorporated into the RdRP database for another round of virus detection. This procedure was repeated ten times. The resulting hits (21,747,015 in total) were compared to the similarity-based clusters for the removal of false positives and virus classification. The false positives and non-RdRP viral proteins that did not contain a RdRP domain were removed by a HMMs-based search using an updated HMMs profile derived from each RdRP cluster and built as described above. The cut-off was set to score≥40 and the aligned fraction to ≥0.4 to remove false positives and partial virus RdRPs (Figures S1C and S2I).

Finally, the expanded clusters were merged into virus supergroups using a hierarchical method employing the Girvan-Newman algorithm,^62^ with the edge betweenness determined based on median *e*-value threshold of 1*E*-3 for each pair of clusters (Figures S1D, S2J-S2K). Briefly, the merging of clusters employed the following steps: (i) the betweenness of all edges (median *e*-value between clusters) in the network was calculated; (ii) the edge(s) with the highest betweenness were removed; (iii) the betweenness of all edges affected by the removal was recalculated; (iv) steps ii and iii were repeated until no edges remained. All processes related to merging were performed using igraph package v1.3.5^63^ implemented in *R*.

### Benchmarking LucaProt and comparisons to Diamond, HMMscan, HH-suite and PalmScan

We assessed the performance of LucaProt (prob≥0.5) relative to other four virus discovery tools: Diamond v0.9.25.126^58^ (with *e*-value threshold of 1*E*-3), HMMscan v3.3.2^59^ (with score≥30 and aligned fraction≥0.7 as referenced from Zayed *et al.,*^6^), HH-suite v3.3.0^64^ (with default parameters), and PalmScan^61^ (with default parameters). This comparison included recall, precision, false positive and false discovery based on the 2.05 million amino acid sequences (length≥300aa) from contigs assembled from the 50 samples collected in this study, employing a unified criterion and utilizing the same well-curated RdRP database (n=5,979).

True positive (TP) virus hits were determined by simultaneously identifying the true viral RdRPs using both aforementioned tools (i.e., LucaProt, Diamond, HMMscan, HH-suite and PalmScan), and subsequently confirming through comparison against the nr protein database (version 2023.01.09) using the Diamond blastp program v0.9.25.126^58^ with an *e*-value threshold of 1*E*-3. True negative (TN) hits were defined as the subset of sequences recognized as cellular proteins and/or with reads from DNA sequencing results mapping against the corresponding contigs, among a data set comprising 2.05 million sequences. A data set of 1,029,342 true negative sequences was obtained through both approaches. Recall was calculated as the ratio of correctly predicted true positives to the total number of true positives (recall rate=TP*100%/total TP). Precision was calculated as the proportion of all predictions that were correctly predicted as true positives (precision rate=TP*100%/total predictions). The false positive (FP) rate was calculated as the proportion of all true negatives that were incorrectly predicted as positives (false positive rate=FP*100%/total TN). False discovery (FD) was calculated as the proportion of all predictions that were incorrectly predicted as positives (false discovery rate=FP*100%/total predictions).

To compare the computational time required by LucaProt to that of HH-suit, Diamond, HMMscan, and PalmScan, we categorized the test data sets into six groups based on amino acid sequence length: (300, 500), (500, 800), (800, 1,000), (1,000, 3,000), (3,000, 5,000), and >5,000. Each group comprised 50 randomly sampled viral RdRPs and 50 non-viral sequences. A single sequence was used as input to calculate the average time on the same machine of the Alibaba Cloud Elastic Compute Service (ECS). The CPU execution environment comprised 96 cores (vCPU) and 192 GiB memory (instance type: ecs.c8i.24xlarge). For LucaProt, we also recorded the time spent on a single NVIDIA A100 80G GPU (instance type: ecs.gn7e-c16g1.16xlarge).

All scripts of the tools used, containing exact parameters and the database utilized in the benchmarking, are available at https://doi.org/10.6084/m9.figshare.26298802.v14.

### Virus verification

To determine whether the newly discovered viral RdRPs belonged to RNA viruses rather than organisms with DNA genomes we performed three verification experiments. First, the RdRPs were compared against the NCBI nr protein database (version 2023.01.09) using Diamond blastp v0.9.25.126,^58^ with the *e*-value threshold set to 1*E*-3. Proteins with similarity to cellular proteins were removed. In addition, we confirmed the presence of the key RdRP motifs (i.e., the A, B and C motifs) through alignment to all RNA supergroups that possessed these motifs (i.e. motif A [DxxxxD], motif B [(S/T)Gxxx(T/G)xxxN], motif C [(S/G/N)DD]). Sequences that failed to cover these motifs were removed.

Second, to ensure the accuracy of DNA read mapping, an initial quality control step was performed on viral contigs using bbduk.sh (https://sourceforge.net/projects/bbmap/). As well as the simultaneous RNA and DNA extraction and sequencing of the 50 environmental samples collected in this study, we searched for published RNA studies that performed both RNA and DNA sequencing on the samples. To confirm that the detected contigs represented *bona fide* RNA viral genomes, reads from the DNA sequencing data were mapped against the viral contigs using Bowtie2 v2.4.2^65^ with the “end-to-end” setting.

Finally, RT-PCR assays were conducted to validate the presence of RNA organisms within the viral supergroups identified. Two pairs of validation primers were designed for each of the representative RdRP sequences from 17 of the 115 RNA viral supergroups involved in 50 samples collected in this study. These comprised: (i) two known supergroups defined by the ICTV: Astro-Poty, Bunya-Arena; (ii) ten unclassified supergroups derived from this study and other studies: Supergroup022, Supergroup034, Supergroup038, Supergroup053, Supergroup055, Supergroup063, Supergroup086, Supergroup104, Supergroup124, Supergroup180; and (iii) five new supergroups identified here: Supergroup102, Supergroup167, Supergroup175, Supergroup184, Supergroup187. We also included gene sequences from two DNA virus families (the *Podoviridae* and *Siphoviridae*), and RT sequences identified in this study, with a product length of 300-550 bp. For each of these samples, both the reverse-transcribed RNA and the matching DNA underwent simultaneous PCR amplification, and the amplification products were subject to electrophoresis using a 1% agarose gel with GelRed dye, which was subsequently visualized under UV.

### Structural prediction and comparisons between viral RdRPs and homologous proteins

Three-dimensional structures of newly identified viral RdRPs from diverse RNA viral supergroups were predicted from primary sequences using AlphaFold2 v2.3^20^ and visualized using the PyMol software v2.5.4 (http://www.pymol.org/pymol). The pLDDT (predicted local distance difference test) score was measured for each structure prediction as its per-residue estimate of the prediction confidence on a scale from 0-100. Considering the computational resource limitations with such a large data set, ten representative RdRPs from each supergroup were predicted: more than 52.03% of the structures predicted had >70% accuracy, showing that AlphaFold2 was a relatively reliable source of structural information. The previously resolved or predicted structures of viral RdRPs, eukaryotic RdRPs, eukaryotic DdRPs and RTs were compared using the Super algorithm.^66^ Considering that the protein structures have similar molecular weights but substantial conformational variation, the “number of aligned atoms after refinement” option was employed to evaluate the similarity between each pair of proteins. Subsequently, NetworkX (https://networkx.org/) was employed to construct a three-dimensional structure diagram using the “edge-weighted spring embedded” approach, with results then mapped as a scatter plot (depicted in the Figure 4D). Simultaneously, we visualized four viral RdRP domain proteins using PyMol.

### Annotation and characterization of virus genomes

Potential open reading frames (ORFs) of every virus genome were predicted based on two criteria: (i) the predicted amino acid sequences were longer than 200 amino acids in length, and (ii) they were not completely nested within larger ORFs. The annotation of non-RdRP ORFs was mainly based on comparisons of predicted proteins to hidden Markov models (HMMs) collected from the Pfam database (version 2022.03.23, https://pfam-legacy.xfam.org/) using HMMscan v3.3.2 implemented in HMMER (*e*=10, score≥10).^59^ For the remaining ORFs, the annotation was performed by blastp comparisons against the nr protein database with an *e*-value threshold of 1*E*-3.

### Analyses of virome diversity, evolution and ecology

To unveil the diversity of the RNA viruses identified and establish putative novel viral species, we employed a protein sequence identity threshold of 90% and a minimum aligned fraction of 30% for all viral RdRP domains identified in this study as implemented by CD-HIT v4.8.1 (https://github.com/weizhongli/cdhit). Abundance levels were subsequently estimated for every putative virus species based on the number of non-rRNA reads per million (RPM) in each sample (i.e., sequencing run) mapped to the RdRP sequences of all viral species using Bowtie2 v2.4.2^65^ with the “end-to-end” setting. The abundance of each putative virus species was estimated as the number of mapped reads per million total non-rRNA reads (RPM) in each library.

Virus alpha diversity (measured using the Shannon index) and overall abundance were subsequently estimated and compared across different geographic locations and ecosystem subtypes, namely; soil, marine, freshwater, wetland, hot spring, salt marsh, and other subtypes. “Marker virus species”, that were greatly enriched in certain ecosystem subtypes, were also identified based on the virus mapping results. The marker virus species were defined as those present only in one ecosystem subtype with RPM≥1 and coverage (percentage of horizontal covered bases with depth≥1) set at ≥20%.

To reveal the diversity and evolutionary relationships of RNA viruses within individual virus supergroups, all previously documented RNA viruses were incorporated into phylogenetic analyses, including RefSeq RdRPs sequences as well as those from previous studies. For supergroups containing more than 500 RdRPs, similarity-based clustering using CD-HIT v4.8.1 (https://github.com/weizhongli/cdhit) with a threshold of identity=0.6 was employed to select representative RdRPs to ensure phylogenetic diversity, while for those containing fewer than 500 RdRPs, all sequences were used in the phylogenetic analysis. The RdRPs of each supergroup were aligned using the L-INS-I algorithm implemented in Mafft v7.475.^60^ Phylogenetic analyses were performed based on the sequence alignment using a maximum likelihood algorithm, employing the LG model of amino acid substitution, a Subtree Pruning and Regrafting (SPR) branch swapping algorithm, and a Shimodaira– Hasegawa-like procedure as implemented in the PhyML program v3.1.^67^

## QUANTIFICATION AND STATISTICAL ANALYSIS

All computational and statistical analyses were performed with the open-source software tools referenced in the STAR Methods along with the described procedures.

## Supplemental video and Excel table titles and legends

**Table S1.** Detailed information on the 10,437 metatranscriptomics retrieved from the SRA database, related to Figure 1 and 2.

**Table S2.** Detailed information on the 50 environmental samples collected in this study, related to Figure 1 and 2.

**Table S3.** Information on the RdRP sequences identified in this study, related to Figure 1.

**Table S4.** Predicted results of RdRPs identified in this study using three methods (Threshold: BLAST: *e*≤1E-3; HMM: score≥10; AI: prob≥0.5), related to Figure 4.

**Table S5.** Viral supergroups identified in 50 collected samples were confirmed through DNA read mapping and RT-PCR, related to Figure 4.

**Table S6.** Size information (the number of dedup-contigs) of all viral supergroups, related to Figure 5.

**Table S7.** Normalized abundance levels (measured by RPM) of each viral species in environmental samples (RPM≥1, coverage≥20%), related to Figure 7.

## REFERENCES

1. Shi, M., Lin, X.D., Tian, J.H., Chen, L.J., Chen, X., Li, C.X., Qin, X.C., Li, J., Cao, J.P., Eden, J.S., et al. (2016). Redefining the invertebrate RNA virosphere. Nature 540, 539–543. 10.1038/nature20167.

2. Shi, M., Lin, X.D., Chen, X., Tian, J.H., Chen, L.J., Li, K., Wang, W., Eden, J.S., Shen, J.J., Liu, L., et al. (2018). The evolutionary history of vertebrate RNA viruses. Nature 556, 197–202. 10.1038/s41586-018-0012-7.

3. Rivarez, M.P.S., Pecman, A., Bačnik, K., Maksimović, O., Vučurović, A., Seljak, G., Mehle, N., Gutiérrez-Aguirre, I., Ravnikar, M., and Kutnjak, D. (2023). In-depth study of tomato and weed viromes reveals undiscovered plant virus diversity in an agroecosystem. Microbiome 11, 60. 10.1186/s40168-023-01500-6.

4. Sutela, S., Forgia, M., Vainio, E.J., Chiapello, M., Daghino, S., Vallino, M., Martino, E., Girlanda, M., Perotto, S., and Turina, M. (2020). The virome from a collection of endomycorrhizal fungi reveals new viral taxa with unprecedented genome organization. Virus Evolution 6, veaa076. 10.1093/ve/veaa076.

5. Wolf, Y.I., Silas, S., Wang, Y., Wu, S., Bocek, M., Kazlauskas, D., Krupovic, M., Fire, A., Dolja, V.V., and Koonin, E.V. (2020). Doubling of the known set of RNA viruses by metagenomic analysis of an aquatic virome. Nat Microbiol 5, 1262–1270. 10.1038/s41564-020-0755-4.

6. Zayed, A.A., Wainaina, J.M., Dominguez-Huerta, G., Pelletier, E., Guo, J., Mohssen, M., Tian, F., Pratama, A.A., Bolduc, B., Zablocki, O., et al. (2022). Cryptic and abundant marine viruses at the evolutionary origins of Earth’s RNA virome. Science 376, 156–162. 10.1126/science.abm5847.

7. Chen, Y.M., Sadiq, S., Tian, J.H., Chen, X., Lin, X.D., Shen, J.J., Chen, H., Hao, Z.Y., Wille, M., Zhou, Z.C., et al. (2022). RNA viromes from terrestrial sites across China expand environmental viral diversity. Nat Microbiol 7, 1312–1323. 10.1038/s41564-022-01180-2.

8. Edgar, R.C., Taylor, J., Lin, V., Altman, T., Barbera, P., Meleshko, D., Lohr, D., Novakovsky, G., Buchfink, B., Al-Shayeb, B., et al. (2022). Petabase-scale sequence alignment catalyses viral discovery. Nature 602, 142–147. 10.1038/s41586-021-04332-2.

9. Obbard, D.J., Shi, M., Roberts, K.E., Longdon, B., and Dennis, A.B. (2020). A new lineage of segmented RNA viruses infecting animals. Virus Evol 6, vez061. 10.1093/ve/vez061.

10. Neri, U., Wolf, Y.I., Roux, S., Camargo, A.P., Lee, B., Kazlauskas, D., Chen, I.M., Ivanova, N., Zeigler Allen, L., Paez-Espino, D., et al. (2022). Expansion of the global RNA virome reveals diverse clades of bacteriophages. Cell 185, 4023–4037 e4018. 10.1016/j.cell.2022.08.023.

11. Urayama, S.I., Fukudome, A., Hirai, M., Okumura, T., Nishimura, Y., Takaki, Y., Kurosawa, N., Koonin, E.V., Krupovic, M., and Nunoura, T. (2024). Double-stranded RNA sequencing reveals distinct riboviruses associated with thermoacidophilic bacteria from hot springs in Japan. Nat Microbiol 9, 514–523. 10.1038/s41564-023-01579-5.

12. Lee, B.D., Neri, U., Roux, S., Wolf, Y.I., Camargo, A.P., Krupovic, M., Consortium, R.N.A.V.D., Simmonds, P., Kyrpides, N., Gophna, U., et al. (2023). Mining metatranscriptomes reveals a vast world of viroid-like circular RNAs. Cell 186, 646–661 e644. 10.1016/j.cell.2022.12.039.

13. Forgia, M., Navarro, B., Daghino, S., Cervera, A., Gisel, A., Perotto, S., Aghayeva, D.N., Akinyuwa, M.F., Gobbi, E., Zheludev, I.N., et al. (2023). Hybrids of RNA viruses and viroid-like elements replicate in fungi. Nat Commun 14, 2591. 10.1038/s41467-023-38301-2.

14. Zheludev, I.N., Edgar, R.C., Lopez-Galiano, M.J., de la Pena, M., Babaian, A., Bhatt, A.S., and Fire, A.Z. (2024). Viroid-like colonists of human microbiomes. bioRxiv. 10.1101/2024.01.20.576352.

15. Dominguez-Huerta, G., Wainaina, J.M., Zayed, A.A., Culley, A.I., Kuhn, J.H., and Sullivan, M.B. (2023). The RNA virosphere: How big and diverse is it? Environ Microbiol 25, 209–215. 10.1111/1462-2920.16312.

16. Cobbin, J.C., Charon, J., Harvey, E., Holmes, E.C., and Mahar, J.E. (2021). Current challenges to virus discovery by meta-transcriptomics. Curr Opin Virol 51, 48–55. 10.1016/j.coviro.2021.09.007.

17. McNutt, A.T., Francoeur, P., Aggarwal, R., Masuda, T., Meli, R., Ragoza, M., Sunseri, J., and Koes, D.R. (2021). GNINA 1.0: molecular docking with deep learning. J Cheminform 13, 43. 10.1186/s13321-021-00522-2.

18. Pham, T.H., Qiu, Y., Zeng, J., Xie, L., and Zhang, P. (2021). A deep learning framework for high-throughput mechanism-driven phenotype compound screening and its application to COVID-19 drug repurposing. Nat Mach Intell 3, 247–257. 10.1038/s42256-020-00285-9.

19. Du, B.-X., Qin, Y., Jiang, Y.-F., Xu, Y., Yiu, S.-M., Yu, H., and Shi, J.-Y. (2022). Compound-protein interaction prediction by deep learning: Databases, descriptors and models. Drug Discov Today 27, 1350–1366. 10.1016/j.drudis.2022.02.023.

20. Jumper, J., Evans, R., Pritzel, A., Green, T., Figurnov, M., Ronneberger, O., Tunyasuvunakool, K., Bates, R., Zidek, A., Potapenko, A., et al. (2021). Highly accurate protein structure prediction with AlphaFold. Nature 596, 583–589. 10.1038/s41586-021-03819-2.

21. Gligorijevic, V., Renfrew, P.D., Kosciolek, T., Leman, J.K., Berenberg, D., Vatanen, T., Chandler, C., Taylor, B.C., Fisk, I.M., Vlamakis, H., et al. (2021). Structure-based protein function prediction using graph convolutional networks. Nat Commun 12, 3168. 10.1038/s41467-021-23303-9.

22. Xu, L., Magar, R., and Barati Farimani, A. (2022). Forecasting COVID-19 new cases using deep learning methods. Comput Biol Med 144, 105342. 10.1016/j.compbiomed.2022.105342.

23. Deng, L. (2014). Deep Learning: Methods and Applications. Foundations and Trends® in Signal Processing 7, 197–387. 10.1561/2000000039.

24. Sarker, I.H. (2021). Deep Learning: A Comprehensive Overview on Techniques, Taxonomy, Applications and Research Directions. SN Comput Sci 2, 420. 10.1007/s42979-021-00815-1.

25. Shang, J., and Sun, Y. (2021). CHEER: HierarCHical taxonomic classification for viral mEtagEnomic data via deep leaRning. Methods 189, 95–103. 10.1016/j.ymeth.2020.05.018.

26. Sukhorukov, G., Khalili, M., Gascuel, O., Candresse, T., Marais-Colombel, A., and Nikolski, M. (2022). VirHunter: A Deep Learning-Based Method for Detection of Novel RNA Viruses in Plant Sequencing Data. Front Bioinform 2, 867111. 10.3389/fbinf.2022.867111.

27. Miao, Y., Liu, F., Hou, T., and Liu, Y. (2022). Virtifier: a deep learning-based identifier for viral sequences from metagenomes. Bioinformatics 38, 1216–1222. 10.1093/bioinformatics/btab845.

28. Liu, F., Miao, Y., Liu, Y., and Hou, T. (2022). RNN-VirSeeker: A Deep Learning Method for Identification of Short Viral Sequences From Metagenomes. IEEE/ACM Trans Comput Biol Bioinform 19, 1840–1849. 10.1109/TCBB.2020.3044575.

29. Lecun, Y., Bottou, L., Bengio, Y., and Haffner, P. (1998). Gradient-based learning applied to document recognition. Proceedings of the IEEE 86, 2278–2324. 10.1109/5.726791.

30. Jordan, M.I. (1997). Serial Order: A Parallel Distributed Processing Approach. 121, 471–495.

31. Vaswani, A., Shazeer, N., Parmar, N., Uszkoreit, J., Jones, L., Gomez, A.N., Kaiser, L., and Polosukhin, I. (2017). Attention Is All You Need. arXiv.

32. Kabir, A., and Shehu, A. (2022). GOProFormer: A Multi-Modal Transformer Method for Gene Ontology Protein Function Prediction. Biomolecules 12. 10.3390/biom12111709.

33. Cao, Y., and Shen, Y. (2021). TALE: Transformer-based protein function Annotation with joint sequence–Label Embedding. Bioinformatics 37, 2825–2833. 10.1093/bioinformatics/btab198.

34. Nambiar, A., Liu, S., Hopkins, M., Heflin, M., Maslov, S., and Ritz, A. (2020). Transforming the Language of Life: Transformer Neural Networks for Protein Prediction Tasks. bioRxiv.

35. Olendraite, I., Brown, K., and Firth, A.E. (2023). Identification of RNA Virus-Derived RdRp Sequences in Publicly Available Transcriptomic Data Sets. Mol Biol Evol 40. 10.1093/molbev/msad060.

36. Avsec, Ž., Agarwal, V., Visentin, D., Ledsam, J.R., Grabska-Barwinska, A., Taylor, K.R., Assael, Y., Jumper, J., Kohli, P., and Kelley, D.R. (2021). Effective gene expression prediction from sequence by integrating long-range interactions. Nature Methods 18, 1196–1203. 10.1038/s41592-021-01252-x.

37. Neuman, B.W., Smart, A., Vaas, J., Bartenschlager, R., Seitz, S., Gorbalenya, A.E., Caliskan, N., and Lauber, C (2024). RNA genome expansion up to 64 kb in nidoviruses is host constrained and associated with new modes of replicase expression. bioRxiv 2024.07.07.602380; doi: 10.1101/2024.07.07.602380.

38. Felipe Benites, L., Stephens, T.G., Van Etten, J., James, T., Christian, W.C., Barry, K., Grigoriev, I.V., McDermott, T.R., and Bhattacharya, D. (2024). Hot springs viruses at Yellowstone National Park have ancient origins and are adapted to thermophilic hosts. Communications Biology 7, 312. 10.1038/s42003-024-05931-1.

39. Thomas, E., Anderson, R.E., Li, V., Rogan, L.J., and Huber, J.A. (2021). Diverse Viruses in Deep-Sea Hydrothermal Vent Fluids Have Restricted Dispersal across Ocean Basins. mSystems 6, e0006821. 10.1128/mSystems.00068-21.

40. Krishnamurthy, S.R., and Wang, D. (2017). Origins and challenges of viral dark matter. Virus Res 239, 136–142. 10.1016/j.virusres.2017.02.002.

41. Chen, J., Guo, M., Wang, X., and Liu, B. (2016). A comprehensive review and comparison of different computational methods for protein remote homology detection. Brief Bioinform 19, 231–244. 10.1093/bib/bbw108.

42. Monttinen, H.A.M., Ravantti, J.J., and Poranen, M.M. (2021). Structure Unveils Relationships between RNA Virus Polymerases. Viruses 13. 10.3390/v13020313.

43. Wolf, Y.I., Kazlauskas, D., Iranzo, J., Lucia-Sanz, A., Kuhn, J.H., Krupovic, M., Dolja, V.V., and Koonin, E.V. (2018). Origins and Evolution of the Global RNA Virome. mBio 9. 10.1128/mBio.02329- 18.

44. Koonin, E.V., Dolja, V.V., Krupovic, M., Varsani, A., Wolf, Y.I., Yutin, N., Zerbini, F.M., and Kuhn, J.H. (2020). Global Organization and Proposed Megataxonomy of the Virus World. Microbiol Mol Biol Revm 84. 10.1128/MMBR.00061-19.

45. Wu, R., Bottos, E.M., Danna, V.G., Stegen, J.C., Jansson, J.K., and Davison, M.R. (2022). RNA Viruses Linked to Eukaryotic Hosts in Thawed Permafrost. mSystems 7, e0058222. 10.1128/msystems.00582-22.

46. Charon, J., Murray, S., and Holmes, E.C. (2021). Revealing RNA virus diversity and evolution in unicellular algae transcriptomes. Virus Evolution 7, veab070. 10.1093/ve/veab070.

47. Ibarbalz, F.M., Henry, N., Brandao, M.C., Martini, S., Busseni, G., Byrne, H., Coelho, L.P., Endo, H., Gasol, J.M., Gregory, A.C., et al. (2019). Global Trends in Marine Plankton Diversity across Kingdoms of Life. Cell 179, 1084–1097 e1021. 10.1016/j.cell.2019.10.008.

48. Kalu, E.I., Reyes-Prieto, A., and Barbeau, M.A. (2023). Community dynamics of microbial eukaryotes in intertidal mudflats in the hypertidal Bay of Fundy. ISME Communications 3, 21. 10.1038/s43705-023-00226-8.

49. Bollback, J.P., and Huelsenbeck, J.P. (2001). Phylogeny, genome evolution, and host specificity of single-stranded RNA bacteriophage (family Leviviridae). J Mol Evol 52, 117–128.

50. Poranen, M.M., Mäntynen, S., and Ictv Report, C. (2017). ICTV Virus Taxonomy Profile: Cystoviridae. J Gen Virol 98, 2423–2424. 10.1099/jgv.0.000928.

51. Callanan, J., Stockdale, S.R., Shkoporov, A., Draper, L.A., Ross, R.P., and Hill, C. (2018). RNA Phage Biology in a Metagenomic Era. Viruses 10. 10.3390/v10070386.

52. Gan, T., and Wang, D. (2023). Picobirnaviruses encode proteins that are functional bacterial lysins. Proc Natl Acad Sci U S A 120, e2309647120. 10.1073/pnas.2309647120.

53. Sharp, P.M., and Simmonds, P. (2011). Evaluating the evidence for virus/host co-evolution. Curr Opin Virol 1, 436–441. 10.1016/j.coviro.2011.10.018.

54. Li, D., Liu, C.-M., Luo, R., Sadakane, K., and Lam, T.-W. (2015). MEGAHIT: an ultra-fast single-node solution for large and complex metagenomics assembly via succinct de Bruijn graph. Bioinformatics 31, 1674–1676. 10.1093/bioinformatics/btv033.

55. Gage, P. (1994). A New Algorithm for Data Compression. C Users J 12, 23–38. 10.5555/177910.177914.

56. Lin, Z., Akin, H., Rao, R., Hie, B., Zhu, Z., Lu, W., Smetanin, N., Verkuil, R., Kabeli, O., Shmueli, Y., et al. (2023). Evolutionary-scale prediction of atomic-level protein structure with a language model. Science 379, 1123–1130. 10.1126/science.ade2574.

57. He, Y., Wang, C., Zhang, S., Li, N., Li, Z.R., and Zeng, Z.Y. (2022). KG-MTT-BERT: Knowledge Graph Enhanced BERT for Multi-Type Medical Text Classification. arXiv.

58. Buchfink, B., Reuter, K., and Drost, H.-G. (2021). Sensitive protein alignments at tree-of-life scale using DIAMOND. Nat Methods 18, 366–368. 10.1038/s41592-021-01101-x.

59. Potter, S.C., Luciani, A., Eddy, S.R., Park, Y., Lopez, R., and Finn, R.D. (2018). HMMER web server: 2018 update. Nucleic Acids Research 46, W200–W204. 10.1093/nar/gky448.

60. Katoh, K., and Standley, D.M. (2013). MAFFT multiple sequence alignment software version 7: improvements in performance and usability. Mol Biol Evol 30, 772–780. 10.1093/molbev/mst010.

61. Babaian, A., and Edgar, R. (2022). Ribovirus classification by a polymerase barcode sequence. PeerJ 10, e14055. 10.7717/peerj.14055.

62. Girvan, M., and Newman, M.E.J. (2002). Community structure in social and biological networks. Proc Natl Acad Sci U S A 99, 7821–7826.

63. Csárdi, G., and Nepusz, T. (2006). The igraph software package for complex network research.

64. Steinegger, M., Meier, M., Mirdita, M., Vohringer, H., Haunsberger, S.J., and Soding, J. (2019). HH-suite3 for fast remote homology detection and deep protein annotation. BMC Bioinformatics 20, 473. 10.1186/s12859-019-3019-7.

65. Langmead, B., and Salzberg, S.L. (2012). Fast gapped-read alignment with Bowtie 2. Nat Methods 9, 357–359. 10.1038/nmeth.1923.

66. Hasegawa, H., and Holm, L. (2009). Advances and pitfalls of protein structural alignment. Curr Opin Struct Biol 19, 341–348. 10.1016/j.sbi.2009.04.003.

67. Guindon, S., and Gascuel, O. (2003). A simple, fast, and accurate algorithm to estimate large phylogenies by maximum likelihood. Syst Biol 52, 696–704.

68. Liu, J.F., H.Q., Liu, Z.Y, Wang, S., Zhang, Q., and He, Z.M. (2022). A Density-Based Spatial Clustering of Application with Noise Algorithm and its Empirical Research. Highlights in Science, Engineering and Technology 7, 174–179.

